# VPS13B is localized at the cis-trans Golgi complex interface and is a functional partner of FAM177A1

**DOI:** 10.1101/2023.12.18.572081

**Authors:** Berrak Ugur, Florian Schueder, Jimann Shin, Michael G. Hanna, Yumei Wu, Marianna Leonzino, Maohan Su, Anthony R. McAdow, Catherine Wilson, John Postlethwait, Lilianna Solnica-Krezel, Joerg Bewersdorf, Pietro De Camilli

**Affiliations:** Department of Cell Biology, Yale University School of Medicine, New Haven, CT, USA; Department of Neuroscience, Yale University School of Medicine, New Haven, CT, USA; Program in Cellular Neuroscience, Neurodegeneration, and Repair, Yale University School of Medicine, New Haven, CT, USA; Aligning Science Across Parkinson’s (ASAP) Collaborative Research Network, Chevy Chase, MD, USA; HHMI, Yale University School of Medicine, New Haven, CT, USA; Department of Microbial Pathogenesis, Yale University School of Medicine, New Haven, CT, USA; Department of Developmental Biology, Washington University School of Medicine, St Louis, MO, USA; Institute of Neuroscience, University of Oregon, Eugene, OR, USA; Nanobiology Institute, Yale University, West Haven, CT, USA; Department of Biomedical Engineering, Yale University, New Haven, CT, USA; Department of Physics, Yale University, New Haven, CT, USA

## Abstract

Mutations in VPS13B, a member of a protein family implicated in bulk lipid transport between adjacent membranes, cause Cohen syndrome. VPS13B is known to be concentrated in the Golgi complex, but its precise location within this organelle and thus the site(s) where it achieves lipid transport remains unclear. Here we show that VPS13B is localized at the interface between cis and trans Golgi sub-compartments and that Golgi complex re-formation after Brefeldin A (BFA) induced disruption is delayed in *VPS13B* KO cells. This delay is phenocopied by loss of FAM177A1, a Golgi complex protein of unknown function reported to be a VPS13B interactor and whose mutations also result in a developmental disorder. In zebrafish, the *vps13b* orthologue, not previously annotated in this organism, genetically interacts with *fam177a1*. Collectively, these findings raise the possibility that bulk lipid transport by VPS13B may play a role in expanding Golgi membranes and that VPS13B may be assisted in this function by FAM177A1.

## INTRODUCTION

Life of eukaryotic cells relies on regulated transport of proteins and lipids from one subcellular compartment to another. Lipid transport is achieved both by membrane traffic as well as by a multitude of lipid transport proteins (LTPs), many of which act at membrane contact sites. Most such proteins function by a piecemeal shuttle mechanism (Saheki and De Camilli, 2017; Balla et al., 2019; Wong et al., 2019; Reinisch and Prinz, 2021). Recently, however, a class of proteins thought to function by a bridge-like mechanism at these sites, and thus optimally suited for the unidirectional bulk delivery of lipids between two closely apposed membranes, has been identified (Levine, 2022; Neuman et al., 2022; Hanna et al., 2023). One such protein is Vps13 (Dziurdzik and Conibear, 2021; Leonzino et al., 2021). While the single yeast Vps13 protein has multiple localizations and functions, the four mammalian VPS13 proteins have distinct, but partially overlapping functions. Mutations in each of them cause severe neurological disorders (Ugur et al., 2020): Chorea-Acanthocytosis (VPS13A) (Rampoldi et al., 2001), Cohen Syndrome (VPS13B) (Kolehmainen et al., 2003), Parkinson’s disease (VPS13C) (Lesage et al., 2016; Darvish et al., 2018; Schormair et al., 2018) and several neurological disorders or embryonic lethality in the case of complete loss-of-function (VPS13D) (Gauthier et al., 2018; Seong et al., 2018; Koh et al., 2020). VPS13A, VPS13C and VPS13D are localized at contacts between the ER and other organelles (Kumar et al., 2018; Yeshaw et al., 2019; Guillen-Samander et al., 2021; Cai et al., 2022; Hancock-Cerutti et al., 2022), while VPS13B is localized predominantly in the Golgi complex (Seifert et al., 2011), with additional pools on lipid droplets (Du et al., 2023) and possibly at endosomes (Koike and Jahn, 2019) but not, at least so far, at membrane contacts with the ER. VPS13B is also the most divergent in amino acid sequence among the 4 paralogues (Levine, 2022).

Consistent with a localization of VPS13B in the Golgi complex (Seifert et al., 2011; Seifert et al., 2015), a less compact Golgi structure and an altered N-glycosylation pattern was observed in cells from Cohen syndrome patients and in several mammalian VPS13B knock-out and knock-down cell lines (Seifert et al., 2011; Duplomb et al., 2014; Zorn et al., 2022). Moreover, Vps13b is required for the growth of the acrosomal membrane of sperm, a Golgi complex-derived membrane critical for fertilization, consistent with the proposed lipid transport role of VPS13 family proteins in membrane expansion, and absence of Vps13b in mice leads to male sterility (Da Costa et al., 2019) and neuroanatomical phenotypes (Montillot et al., 2023). However, the precise localization of VPS13B within the Golgi complex, and thus the sites where its lipid transport function is likely achieved, remains unknown.

Here, we show that VPS13B mainly localizes on the cis side of the Golgi complex and is especially enriched at the interface between cis-Golgi and trans-Golgi membranes. We demonstrate that loss of VPS13B in HeLa cells delays Golgi complex recovery after its Brefeldin A (BFA)-induced dispersion, raising the possibility that VPS13B’s lipid transfer function may have a role in Golgi complex reformation. We further provide evidence in mammalian cells and in zebrafish for a functional partnership between VPS13B and FAM177A1, a recently identified Golgi complex protein (Fasimoye et al., 2023; Hickey et al., 2023)(Legro et al., in revision) of unknown function implicated in a human neurodevelopmental disorder (Alazami et al., 2015)(Legro et al., in revision).

## RESULTS

### Predicted Structure of VPS13B

Comparative analysis of the primary sequences of the four mammalian VPS13 proteins reveals that VPS13B is the most divergent among them (Velayos Baeza et al., 1993; Levine, 2022). Such divergence occurred early in evolution, as in protists and algae, whose genome includes only two VPS13 genes, one of these two genes encodes a protein with similarity to VPS13B, while the other appears to be the ancestor of VPS13A, VPS13C and VPS13D. In spite of the primary sequence divergence, fold prediction algorithms (Yang et al., 2020; Jumper et al., 2021) reveal that VPS13B has all the defining features of VPS13 family proteins: a rod-like core (∼27-nm long) comprising 13RBG motifs, flanked at its C-terminal portion by folded domains with targeting and likely regulatory functions (Fig. 1A, S1A and S1B). One distinct feature of VPS13B is the presence of an accessory folded domain, a module with a Jelly-roll fold (Dall’Armellina et al., 2022; Levine, 2022; Hanna et al., 2023), which is an outpocketing of the RBG rod just upstream of the VAB domain (shown in gray in Fig. S1A, S1B and S1C). VPS13A, VPS13C and VPS13D lack this domain but have instead a WWE domain (VPS13A and VPS13C) and a ricin-B domain (VPS13D) (Hanna et al., 2023).

**Figure 1.**
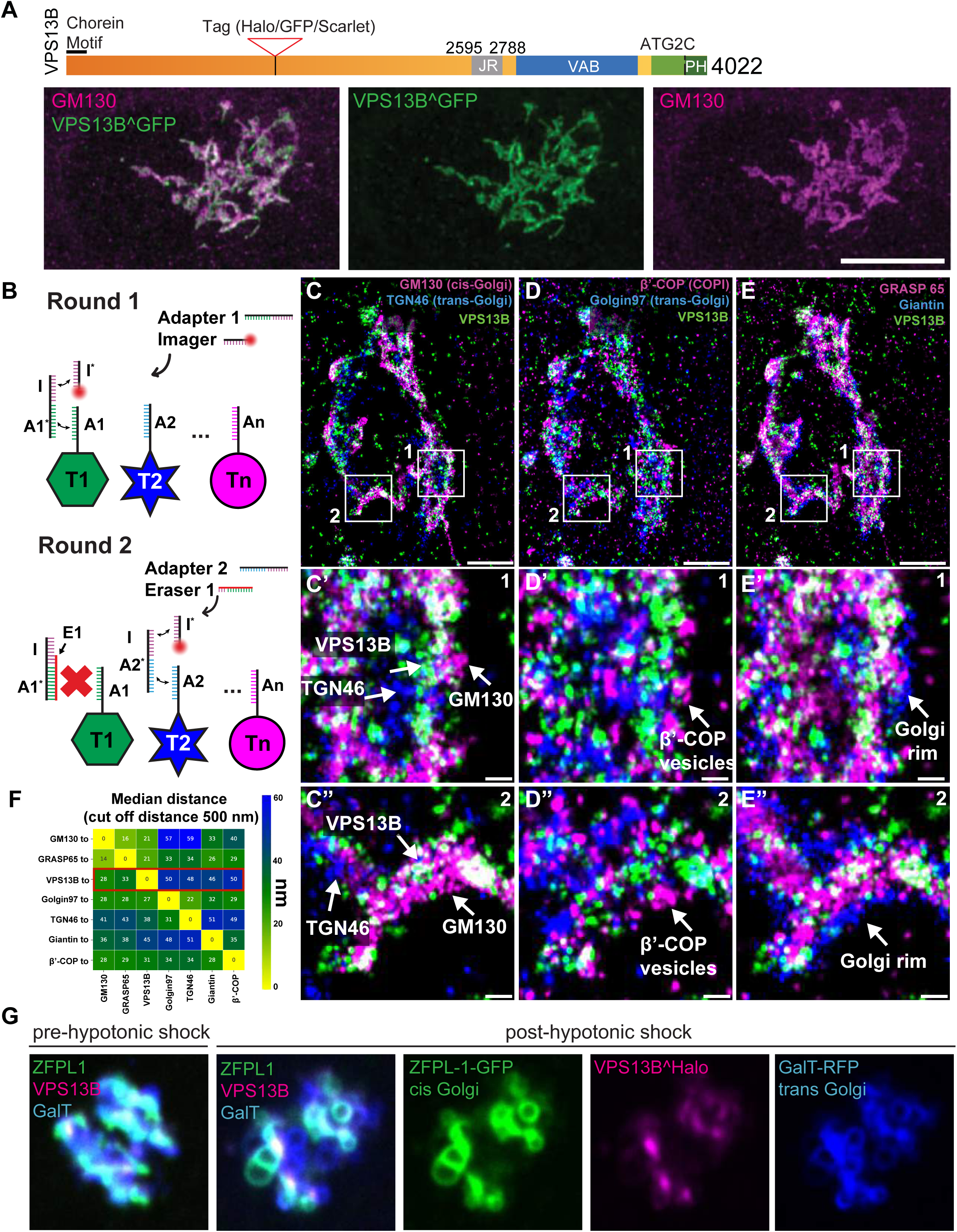
VPS13B is localized at the interface between cis-Golgi and trans-Golgi membranes. **(A)** Domain structure of human VPS13B (upper panel), COS7 cells expressing codon optimized human VPS13B^GFP immunolabeled for GFP and GM130. Scale bar = 10 µm (lower panel) **(B)** Schematic representation of labeling steps used for FLASH-PAINT. FLASH-PAINT performed in HeLa cells expressing VPS13B^GFP and immunolabeled with anti-GFP. **(C-E)** FLASH-PAINT signals of a Golgi complex immunolabeled for the indicated Golgi complex targets. Scale bar = 2 µm. High magnification fields of the boxed areas in fields **(C-E)** are shown below in field **(C’-E’)** and **(C”-E”)**. Scale bar = 250 nm. **(F)** Median distances between super-resolved signals of different targets. Only signals closer than 500 nm to each other were considered. **(G)** Live imaging snapshots COS7 cell expressing VPS13B^halo, ZFPL1-GFP (a cis Golgi marker) and GalT-RFP (a trans-Golgi marker) before and after (8 min) hypotonic shock. Scale bar = 5 µm

*VPS13B is localized at the interface between cis-Golgi and trans-Golgi membranes*.

Due to low levels of endogenous protein expression and inconsistencies with commercially available antibodies for immunocytochemistry, to gain further insight into the subcellular targeting of VPS13B, we expressed codon optimized versions of full-length human VPS13B, - fused to either GFP (VPS13B^GFP) or Halo (VPS13^Halo), in COS7 cells. As we had observed with other VPS13 proteins (Guillen-Samander et al., 2021; Cai et al., 2022), codon optimization helps achieving robust expression of these very large proteins. GFP or Halo were inserted into the unstructured loop that connects the 5^th^ and 6^th^ repeating RBG motifs, a position predicted not to interfere with the hydrophobic channel and thus with lipid transport activity (Fig. S1A and S1D). In agreement with previous reports (Seifert et al., 2011; Seifert et al., 2015), VPS13B^GFP roughly co-localized with immunoreactivity for the cis-Golgi protein GM130 (Fig. 1A). However, the fluorescence patterns of the two proteins did not completely overlap. To gain more precise insight into the localization of VPS13B within the Golgi complex we used two different strategies.

In one strategy, we performed super-resolution FLASH-PAINT imaging (Schueder et al., 2023), a variant of DNA-PAINT (Jungmann et al., 2010; Schnitzbauer et al., 2017) enabling efficient multiplexed imaging. FLASH-PAINT utilizes transient binding of short diffusing fluorescently labeled DNA oligomers (termed ‘imagers’) to single-stranded DNA docking sites at the target (in our case antibodies to specific proteins) via a transient adapter, resulting in apparent fluorescent blinking at these sites (Fig. 1B). In a subsequent analysis, super-resolved coordinates of the docking sites are extracted from the observed distinct single-molecule blinking events.

To enable sequential multiplexed imaging of VPS13B^GFP (using GFP nanobodies) relative to different Golgi complex proteins, seven protein species were each labeled with an antibody/nanobody conjugated to a different single-stranded DNA docking site. For imaging, a transient adapter was used to mediate the interaction between the imager and the docking site. To switch from imaging one target species to the next, a single-stranded DNA eraser (Fig. 1B) was added after completion of a round of imaging. This eraser binds to the adapter from the previous round, preventing the adapter from binding to the docking site and, therefore, effectively eliminating the signal. This procedure was then repeated for all target antigens. An analysis of distances of the various antigens from each other revealed that VPS13B was closest to the cis-Golgi complex markers GM130 (on average ∼28 nm) (Fig. 1C, 1C’ and 1C”) and GRASP65 (on average ∼ 33 nm) (Fig. 1E, 1E’ and 1E”), and the furthest away from trans Golgi markers TGN46 and Golgin97 (Fig. 1C, 1C’, 1C”, 1D, 1D’, 1D” and 1F). VPS13B was also at some distance from β-COP positive signal (Fig. 1D, 1D’, Fig1D” and 1F), indicating that VPS13B is not localized at the interface of the cis-Golgi complex with the intermediate compartment and the ER, but on portions of the cis-Golgi complex distal to such interface.

As an alternative strategy, we exposed live cells to a hypotonic shock. This treatment rapidly expands the volume of the cytoplasm, thus decreasing organelle crowding and causing partial release of cytosolic proteins. It also causes membrane organelles to swell, but, at least within short time ranges (minutes), does not disrupt their membrane contacts, which in fact may coalesce into fewer and larger contacts (King et al., 2020; Guillen-Samander et al., 2021). As a result, the localization of proteins at membrane contact sites can be more easily visualized than in intact cells. In hypotonically treated COS7 cells expressing VPS13B^Halo, ZFPL1-GFP (a cis-Golgi marker (Chiu et al., 2008)) and GalT-RFP (a trans-Golgi marker (Roth and Berger, 1982)) Golgi cisternae underwent rapid swelling resulting in two sets of closely apposed, but distinct vacuoles positive for the cis and for the trans-Golgi marker, respectively (Fig. 1G, Video 1). Interestingly, VPS13B selectively localized at the interface of the two types of vacuoles suggesting that it may function as a tether between cis-and trans-Golgi membranes and consistent with the FLASH-PAINT results (Fig. 1C’). We also performed similar experiments with cells expressing tagged VPS13B and markers of the ER and of the Golgi complex but did not find convincing evidence for presence of VPS13B at the interface of ER and Golgi membranes, where other lipid transport proteins are known to function (De Matteis et al., 2007; Venditti et al., 2020). This is consistent with the lack of an FFAT motif for ER binding in VPS13B, a motif that is present in VPS13A, VPS13C and VPS13D (in the latter case a phosphoFFAT motif (Guillen-Samander et al., 2021)).

While performing the localization studies, we noticed that overexpressed VPS13B can also localize to lipid droplets, more so when the temperature is lowered to 20°C. A previous study indicated that VPS13B can be present at the Golgi complex-lipid droplet interface (Du et al., 2023), although our results indicate that VPS13B can localize to lipid droplets independent of their proximity to Golgi complex (Fig. S1E). Of note, Du *et al*., also report no evidence for a localization of VPS13B at the ER, or at the interface of the ER with other organelles.

Overall, our results suggest that in contrast to other VPS13 proteins, VPS13B may bridge cis and trans Golgi membranes.

### FAM177A1 is a neighbor of VPS13B in the Golgi complex

Previously, the localization of VPS13B at the Golgi complex was reported to be dependent on Rab6 (Seifert et al., 2015), a Rab family member with multiple reported sites of action in the Golgi complex (Feldmann et al., 1995). Further elucidation of the properties and physiological function of VPS13B will be helped by the identification of potential functional partners. These proteins may assist VPS13B in its tethering function or in lipid extraction and delivery from and to membranes. Recently, we have shown that FAM177A1, a protein of unknown function identified as a potential interactor of VPS13B by several high-throughput proteomic studies (Huttlin et al., 2015; Huttlin et al., 2017; Huttlin et al., 2021) (Fig. S2A) is localized in the Golgi complex (Fasimoye et al., 2023)(Legro et al., in revision) (Fig. S2B). Interestingly, recessive loss-of-function mutations in *FAM177A1* are responsible for a neurodevelopmental disorder that displays characteristics similar to those of Cohen syndrome, as reported in (Alazami et al., 2015) and supported by additional still unpublished cases (Legro et al., in revision). This raises the possibility that FAM177A1 may be a functional partner of VPS13B. Based on folding predictions (Jumper et al., 2021; Varadi et al., 2022) FAM177A1 is primarily a disordered protein with two α–helices and a short β-sheet hairpin (Fig. 2A), without transmembrane regions.

**Figure 2.**
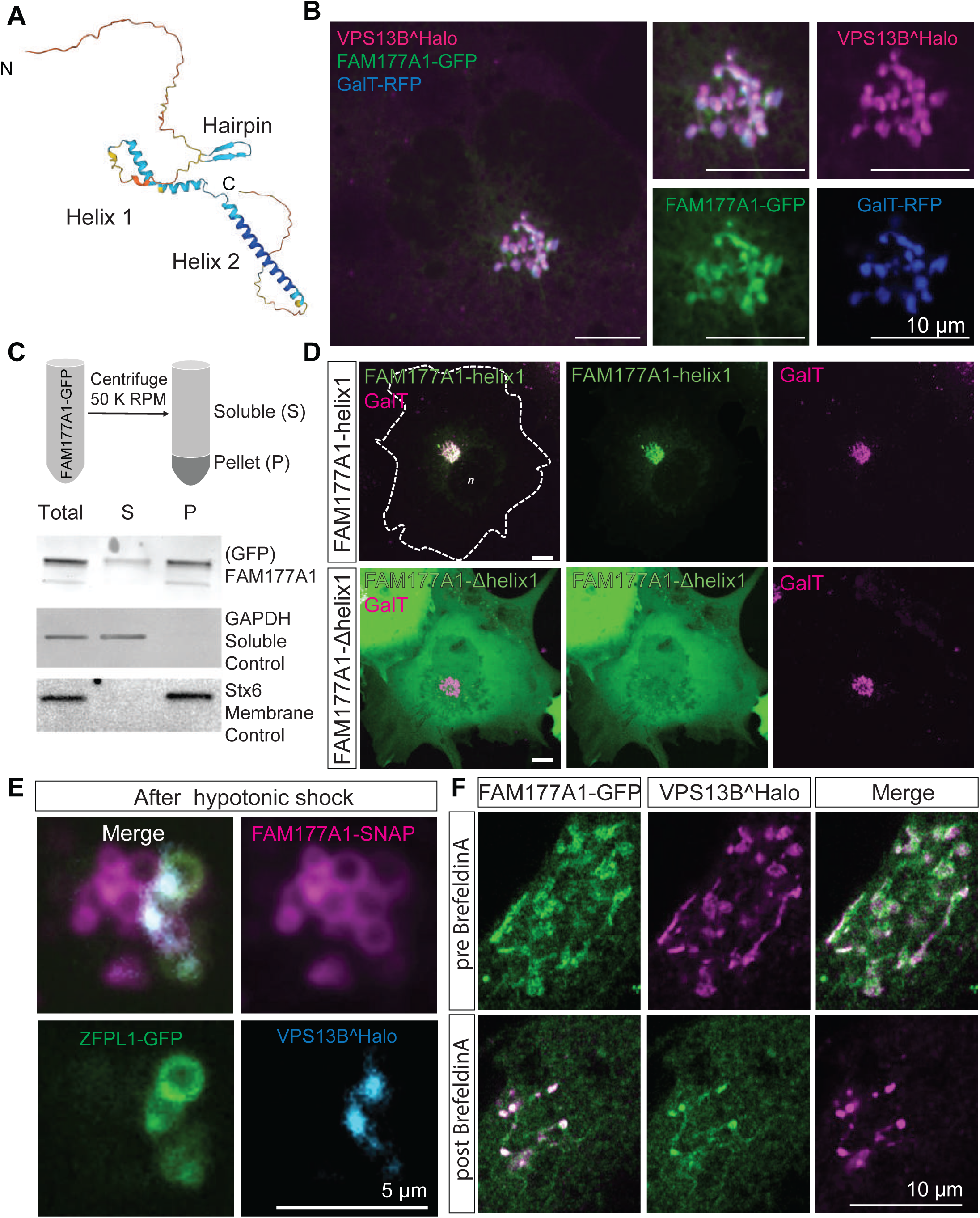
FAM177A1 is a protein neighbor of VPS13B in the Golgi complex. **(A)** Alphafold2 prediction of human FAM177A1 structure. **(B)** COS7 cells expressing VPS13B^Halo, FAM177A1-GFP and GalT-RFP showing that FAM177A1 is localized at the Golgi complex. **(C)** Western blot analysis of soluble cytosolic fraction and total membrane faction of HeLa cells expressing FAM177A1-GFP. GAPDH was used as a soluble control, Stx6 as a membrane control protein. **(D)** Top Panel, co-expression of FAM177A1-helix1-GFP and GalT-RFP in COS7 cells. Bottom Panel, co-expression of FAM177A1-Δhelix 1-GFP and GalT-RFP in COS7 cells. Scale bar = 10 µm. **(E)** Live imaging snapshots of COS7 cells expressing FAM177A1-SNAP, ZFPL1-GFP and GalT-RFP treated with water for 10 minutes. Scale bar = 5 µm**. (F)** Live imaging snapshots of HeLa cells expressing FAM177A1 GFP and VPSB^Halo before and after BFA (5 µg/mL for 50 minutes) treatment. Scale bar = 10 µm.

FAM177A1-GFP (we did not detect any specific localization with the commercially available antibodies) colocalizes with tagged-VPS13B in the Golgi complex (Fig. 2B), although at very high level of expression some signals can also be observed throughout the ER, where VPS13B is never observed. Moreover, we found that its localization in the Golgi complex does not depend on VPS13B, because its localization is preserved in VPS13B KO cells (Fig. S3B). Fractionation of the cell lysate into a cytosolic and a total membrane fraction, followed by western blotting for GFP, revealed that the bulk of FAM177A1-GFP is membrane-associated, but that a small amount remained soluble, while western blotting of the same fractions for a control cytosolic protein (GADPH) and a control intrinsic membrane protein (STX6) showed that they exclusively segregated as expected in the soluble and membrane fractions, respectively (Fig. 2C). These findings confirmed that FAM177A1 is not an intrinsic membrane protein, although it interacts with membranes.

To determine how FAM177A1 is targeted to the Golgi complex, we transfected COS7 cells with constructs encoding the three predicted folded domains of FAM177A1 fused to GFP and assessed their localizations. Helix 1 predominantly localized to the Golgi complex (Fig. 2D, top panel), whereas Helix 2 and the β-sheet hairpin were cytosolic (Fig. S2C). Moreover, a FAM177A1 construct lacking Helix 1 had a diffuse cytosolic localization (Fig. 2D), providing further evidence that Helix 1 is responsible for the Golgi complex localization of FAM177A1.

FAM177A1 has a close paralogue in the human genome, FAM177B (Fig. S2D). Expression of FAM177B-flag in COS7 cells showed that this protein also localizes to the Golgi complex suggesting that the two proteins have overlapping functions (Fig. S2E). Consultation of the Protein Atlas and RNA-seq depositories (https://www.proteinatlas.org/ENSG00000197520-FAM177B) reveals that in contrast to the broad distribution of FAM177A1 across tissues, FAM177B has a more restricted expression and is generally expressed at much lower levels with highest expression reported in the stomach.

To obtain further insight into the localization of FAM177A1, we examined the redistribution of FAM177A1-SNAP in response to hypotonic shock in cells also expressing VPS13B^Halo and the cis-Golgi marker ZFPL1-GFP. After the hypotonic shock, FAM177A1-SNAP decorated both cis- and trans-Golgi complex membranes, whereas ZFPL1-GFP remained confined to the cis-Golgi and VPS13B, as shown in Fig. 1G, was restricted to the cis-trans Golgi interface (Fig. 2E). As the localization of VPS13B was shown to be disrupted by the expression of dominant-negative Rab6a (Rab6a^T27N^) (Seifert et al., 2015), we also investigated whether this Rab6 construct affected the localization of FAM177A1. We found, however that FAM177A1 remained in the Golgi complex upon Rab6a^T27N^ expression (Fig. S2F).

We additionally performed experiments with Brefeldin A (BFA), a fungal metabolite that blocks transport between the ER and Golgi complex by inhibiting Arf1 and induces the reversible disassembly of the Golgi complex (Lippincott-Schwartz et al., 1989). In agreement with its Golgi complex localization, VPS13B was previously shown to disperse by cell treatment with BFA (Duplomb et al., 2014; Seifert et al., 2015) although we found that few spots remained in the region previously occupied by the Golgi complex. When cells co-expressing FAM177A1 and VPS13B were exposed to BFA, both proteins acquired a diffuse distribution, but the remaining VPS13B puncta were now also positive for FAM177A1 (Fig. 2F and Video 2). This result was in contrast to what we observed when cells expressing VPS13B and the Golgi enzyme GalT were treated with BFA. Here, GalT did not colocalize with the remaining VPS13B-positive spots and instead dispersed into the ER as anticipated (Fig. S2G).

### Loss of VPS13B and FAM177A1 leads to delay in Golgi complex re-formation after BFA treatment

It was described previously that loss of VPS13B results in a less compact Golgi structure (Seifert et al., 2011). We generated *VPS13B* KO HeLa cells by CRISPR/Cas9 (Fig. S3A, 3A and 3B) and made similar observations using GM130 as a marker (Fig. 3A). FAM177A1 was still localized in the Golgi in these cells (Fig. S3B), speaking against a role of VPS13B in its recruitment. Given the putative function of VPS13B in the bulk transport of phospholipids between membranes, we explored whether its absence leads to perturbation of Golgi complex dynamics, as assessed by GM130 immunofluorescence. Treatment of VPS13B KO cells with BFA induced dispersion of the Golgi complex with a time course similar to control cells (Fig. 3A). However, Golgi complex reformation was delayed in KO cells relative to control cells upon BFA washout. While the Golgi complex fully reassembled within 2 hours in control cells, its reassembly was still incomplete in about 50% of the cells after 5 hours (Fig. 3A and 3D). This delay was rescued by exogenous expression of VPS13B^GFP.

**Figure 3.**
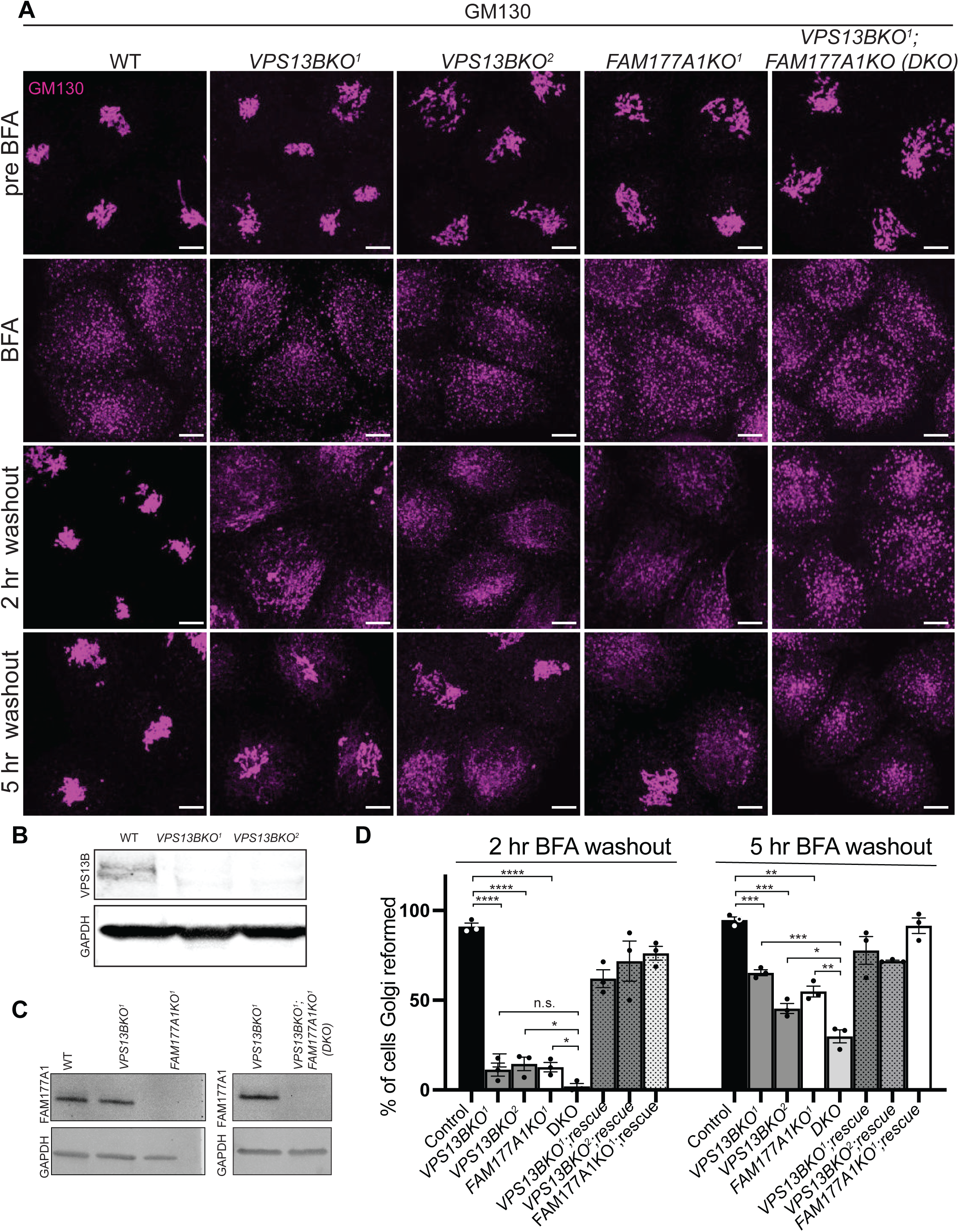
Loss of VPS13B and FAM177A1 leads to delay in Golgi complex reformation after BFA treatment. **(A)** Anti-GM130 immunofluorescence of WT, *VPS13BKO^1^, VPS13BKO^2^*, *FAM177A1KO* and *FAM177A1;VPS13B* double KO cells before incubation with BFA, after 1 hr in BFA (5 µg/mL) and after subsequent washings as indicated; superscripts indicate different clones. **(B and C)** Western blots of total cell homogenates showing loss of the VPS13B band and/or the FAM177A1 band in the KO clones. GAPDH was used as a gel loading control**. (D)** Quantification of Golgi complex reformation in cells with the indicated genotypes after BFA washout for 2 or 5 hrs. “Rescue” refers to exogenous expression of the knocked-out protein. Data are mean ± s.e.m. n = 3 per condition, in each condition 75–100 cells were quantified. Unpaired, two-tailed t-tests. NS, not significant. ***p<0.001; **p<0.01; *p<0.05

We also examined whether loss of *FAM177A1* had an impact on Golgi complex structure and to this aim we generated *FAM177A1* KO HeLa cells (Fig. 3A, 3C and S3C). In these cells the localization of VPS13B was not affected (Fig. S3D) and overall Golgi complex morphology was not severely perturbed with the exception of a less compact structure as assessed by GM130 staining (Fig. 3A). Moreover, dispersion of the Golgi complex upon BFA treatment was not different from controls. However, as in *VPS13B* KO cells, Golgi complex recovery was delayed, and this delay was rescued by exogenous expression of *FAM177A1* (Fig. 3A, 3D).

Finally, we generated *VPS13B* and *FAM177A1* double-KO HeLa cells (DKO, Fig. 3A, 3C). In these cells, Golgi complex recovery following BFA was even more strongly delayed when compared to the recovery in both single KO cells. Following BFA washout, approximately 50% of *FAM177A1* KO cells re-formed their Golgi complex by 5 hours, as shown before for *VPS13B* KO cells (Fig. 3D). However, only approximately 30% of the *VPS13B;FAM177A1* DKO cells displayed recovery of the Golgi complex 5 hours after BFA washout, suggesting a synergistic effect of the two mutations (Fig. 3A, 3D).

### A partnership of fam177a1 and vps13b in zebrafish

To test whether our results in cultured cells had a relevance to organism physiology, we carried out studies in zebrafish. Zebrafish has two FAM177 genes, *fam177a1a* and *fam177a1b*. In contrast, a gene encoding VPS13B ortholog had not been annotated in zebrafish. As a functional partnership between Fam177a1 and Vps13b implies their coexpression, we searched for a *VPS13B* ortholog in the unannotated zebrafish genome sequence by BLAST searches and identified a *vps13b* candidate ortholog. Analysis of its structure revealed strong similarity to mammalian VPS13B, including the presence of a jelly-roll module (Fig. S3E). Furthermore, the candidate *vps13b* gene showed conserved synteny with the human V*PS13B* gene (Fig. S3F). The conservation of sequence and genomic location shows that these two genes are indeed orthologs. In addition, we performed qRT-PCR analysis of *vps13b* transcripts in WT zebrafish embryos at early developmental stages (2-cell, 6 hpf and 24 hpf) and confirmed the expression of *vps13b* at all these stages, with higher levels at 2-cell stage indicating maternal expression (Fig. S3G).

Recently, we had generated large deletions in the two zebrafish *FAM177A1* homologs, *fam177a1a* and *fam177a1b* (*fam177a1a/b*)(Legro et al., in revision). While fish with these mutations, *fam177a1a;fam177a1b Double KO (DKO)* were viable and fertile, they exhibited a delay in reaching WT length during embryogenesis and at 25 hpf were significantly shorter than controls (Fig. 4A, 4B left panels)(Legro et al. in revision). To determine if *fam177a1a/b* functionally interact with zebrafish *vps13b,* we injected WT and *fam177a1a;fam177a1b DKO* embryos at the one-cell stage with two *vps13b* guide RNAs and Cas9 to generate a pool of edited fish (“crispants” (Hwang et al., 2013)) (Fig. S3H). As these injections can only impair zygotic gene function and in a mosaic fashion, the resulting phenotypes would be expected to be a hypomorph phenotype rather than a complete loss-of function phenotype. Crispant fish embryos for *vps13b* displayed regular body length at all stages (Fig. 4A, 4B left panels), whereas *fam177a1a;fam177a1b DKO;vps13b* crispants had significantly shorter (25% reduction compared to WT) body length at 25 hpf when compared to *fam177a1a;fam177a1b DKO* fish at the same stage *(15% reduction compared to WT)* indicating a functional interaction between Fam177a1 proteins and Vps13b (Fig. 4A, 4B left panels). Moreover (compare right and left panels in Fig. 4B), when zebrafish embryos were continuously exposed to BFA from 6 to 25 hpf (0.8 µg/mL, a dose comparable to the concentration used for *in vitro* studies), a treatment that by itself results in a 7% reduction in body length of WT animals at 25 hpf, an additional decrease in body length was observed in *fam177a1a;fam177a1b DKO (30% reduction compared to untreated WT)* and *vps13b* crispants (20% reduction compared to untreated WT) revealing that loss of these proteins sensitizes zebrafish embryos to BFA effects. Although the *fam177a1a;fam177a1b DKO;vps13b* crispants treated with BFA had a significantly decreased body length (35% reduction) when compared to either WT animals or to *vps13b* crispants alone, they did not show a significant decrease when compared to *fam177a1a;fam177a1b DKO* animals. In sum, loss of either *vps13b* or *fam177a1a/b* in zebrafish leads to a BFA sensitivity in animals and the hypomorphic zygotic *vps13b* crispant loss of function enhances *fam177a1a/b* mutant phenotype, suggesting a functional interaction of these two Golgi-localized proteins.

**Figure 4.**
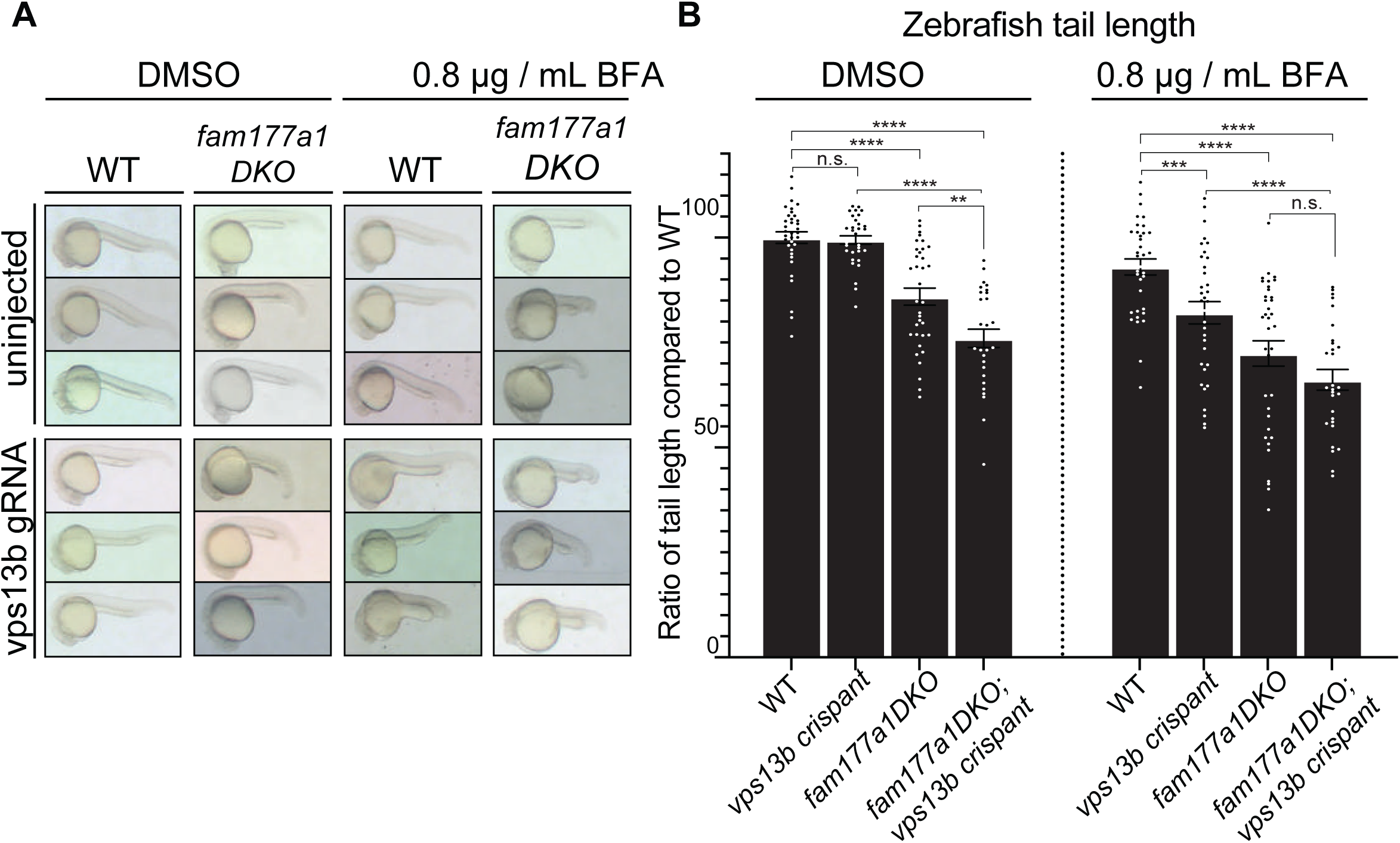
A partnership of *fam177a1* and *vps13b* in zebrafish. **(A)** Representative images of WT and *fam177a1a;fam177a1b double KO (fam177a1DKO)* zebrafish embryos that are either uninjected or injected with *vps13b* gRNAs/Cas9. Left Panel shows embryos treated with DMSO at 6hpf until 25 hpf whereas right panel shows embryos treated with 0.8 µg/mL BFA at 6hpf until 25 hpf. **(B)** Quantifications of tail length results shown in **(A)**. Data are shown as means ± s.e.m., Unpaired, two-tailed t-tests. n.s., not significant. ****p<0.0001; ***p<0.001; **p<0.01.

## DISCUSSION

Our study provides new evidence for the idea that VPS13B differs in some special way from the other three mammalian VPS13 proteins, in agreement with its early evolutionary divergence from the common progenitor of VPS13A, VPS13C and VPS13D (Velayos-Baeza et al., 2004; Levine, 2022). In contrast to what we have found for the other three VPS13 proteins, we did not find evidence for binding of VPS13B to the ER or more specifically at the interface of the ER with other organelles. This negative result is supported by the lack of an FFAT motif in its sequence. Absence of an FFAT motif and lack of evidence for ER binding was also observed for another member of the RBG lipid transport protein family, SHIP164/BLTP3B (Hanna et al., 2022). Of note, the N-terminal regions of both VPS13B and BLTP3B, the so called chorein motifs, have a different overall charge than VPS13A, VPS13C and VPS13D (Levine, 2022). Using FLASH-PAINT, a recently developed super-resolution microscopy technique (Schueder et al., 2023), we have located the protein at the cis side of the Golgi complex, but not at its surface facing the intermediate compartment. Furthermore, we have shown that upon exposure of cells to a hypotonic shock to induce swelling of Golgi complex cisterns, VPS13B coalesces at contacts between large vesicles positive for cis-Golgi and trans-Golgi markers, respectively. Such a localization is consistent with a tethering function of VPS13B, in agreement with what has been shown for other VPS13 family members, but not at an ER contact site. These findings raise the possibility that VPS13B may assist in the transfer of phospholipids from cis-to trans-Golgi membranes via a bridge-like mechanism, and thus operates in parallel to vesicular transport, although this hypothesis remains highly speculative. Our study also shows that lack of VPS13B results in a delay in the re-formation of the Golgi complex after its BFA-induced dispersion. While this finding may have many explanations, a defect in Golgi cisternae reassembly due to a defect in lipid transport between Golgi complex subcompartments is plausible.

A membrane tethering function for VPS13B implies a minimum of two binding sites, one on each of the two membranes connected by this protein. At least one binding site is likely to be Rab6, as VPS13B has been reported to be a Rab6 effector and dominant negative Rab6 was shown to result in a cytosolic localization of VPS13B (Seifert et al., 2015; Du et al., 2023). This is surprising because upon loss of Rab6 binding, VPS13B could be expected to remain associated with the other membranes via its second binding site. It is possible that loss of Rab6 binding may allow the occlusion of the second binding site through an intramolecular interaction. A similar result was obtained for VPS13C, which has binding sites for Rab7 on lysosomes and for VAP on the ER, but becomes diffusely cytosolic upon expression of dominant negative Rab7 (Hancock-Cerutti et al., 2022).

Our study also suggests a functional link of VPS13B to FAM177A1, a hit in proteome-wide screens of VPS13B interactors (Huttlin et al., 2015; Huttlin et al., 2017; Huttlin et al., 2021). Our current study provides strong evidence for a close proximity of pools of the two proteins in the Golgi complex, although, so far, we do not have evidence for a direct interaction between them. FAM177A1 is not needed for the Golgi complex localization of VPS13B and, *vice versa*, FAM177A1 localizes to the Golgi complex even in the absence of VPS13B. Additionally, as shown by the hypotonic lysis experiments, the localization of FAM177A1 in the Golgi complex is more widespread on Golgi membranes than the localization of VPS13B, suggesting that only a small pool of the two proteins may interact with each other at any given time. On the other hand, we have found that the delay in Golgi complex re-formation after BFA-induced dispersion observed in *VPS13B* KO cells is phenocopied in *FAM177A1* KO cells and that the loss of both proteins results in a more severe delay, consistent with a synergistic effect of the two proteins.

Likewise, an additive effect of VPS13B and FAM177A1 is suggested by experiments in zebrafish, where a *vps13b* gene had not been annotated before. In this organism, the developmental defect caused by the absence of *fam177a1a/b* is enhanced by the partial loss of function of *vps13b* (*fam177a1a fam177a1b DKO;vps13b* crispant fish). Moreover, chronic exposure of zebrafish to BFA enhances the developmental phenotypes of *fam177a1DKO*, both in the presence or absence of the additional defect in VPS13B, and unmasks a developmental defect in *vps13b* crispant animals. Most interestingly, loss of either protein in humans result in developmental disorders that include neurological defects. In line with a partnership of VPS13B and FAM177A1, the *C. elegans* genome, which lacks a *VPS13B* gene, also lacks a gene encoding FAM177A1.

In conclusion, our studies provide new insight into the site of action of VPS13B, suggest the possibility that this protein may mediate bridge-like lipid transport between Golgi subcompartments, and identify FAM177A1 as one of its potential functional partners. They provide a foundation for further investigations of the cellular (Seifert et al., 2011; Duplomb et al., 2014; Zorn et al., 2022) and organismal (Da Costa et al., 2019; Kim et al., 2019; Gabrielle et al., 2021; Montillot et al., 2023) phenotypes produced by VPS13B loss-of-function.

## Acknowledgements

We thank Andres-Guillen-Samander for helpful discussion. This work was supported in part by NIH grants R01NS36251 (to PDC), R01OD011116 (to JHP), R01GM151829 (to JB), and R01HD110556 (to LSK), and by the Kavli Institute for Neuroscience (PDC). This research was also funded in part by Aligning Science Across Parkinson’s ASAP-000580 through the Michael J. Fox Foundation for Parkinson’s Research (MJFF). B.U. was a Fellow supported by the William N. & Bernice E. Bumpus Foundation. F.S. acknowledges support from the Human Frontier Science Program (LT000056/2020-C). For the purpose of open access, the author has applied a CC BY public copyright license to all Author Accepted Manuscripts arising from this submission.

## Materials and Methods

### Antibodies and reagents

Primary antibodies used were as follows: rabbit VPS13B (proteintech; 24505-1-AP; RRID:AB_2879579; IB 1:250); rabbit FAM177A1 (Bethyl Laboraties A303-366A; RRID:AB_10952864; IB 1:1000); mouse GM130 (BD Bioscience; 610822; RRID:AB_398141; IF 1:500); rabbit GM130 (Abcam; ab30637; RRID:AB_732675 IF1:300 and proteintech 11308-1-AP); rabbit TGN46 (Abcam; ab50595; RRID:AB_2203289; IF 1:200 and proteintech 10598-1-AP); mouse GAPDH (Proteus; 40-1246; IB 1:10,000); mouse α-Tubulin (Sigma Aldrich; T5168; RRID:AB_477579; IB 1:10000); rabbit GFP (Invitrogen, IF 1:250; IB 1:1000); rabbit Stx6 (Synaptic systems; 110 062; RRID:AB_887854; IB 1:1000); GRASP 65 (Abcam; ab174834; ), Giantin (Sigma, HPA011555) , Golgin 97 (Atlas antibodies; HPA044329); COPI (CMIA10) were customary made in the Rothman lab. Secondary antibodies used were goat anti-mouse IgG (926-32210; LI-COR Biosciences) and goat anti-rabbit IgG (926-68021; LI-COR Biosciences).

Cy3B-modified DNA oligonucleotides were custom-ordered from IDT. DNA labeled Nanobodies were obtained from Massive Photonics. Halo and SNAP tag ligands (JF549, JF646) were kind gifts from L. Lavis (Janelia Research Campus, Ashburn, VA).

Sodium chloride 5 M (cat: AM9759) were obtained from Ambion. Ultrapure water (cat: 10977-015) was purchased from Invitrogen. µ-Slide 8-well chambers (cat: 80807) were purchased from ibidi. Methanol (cat: 9070-05) was purchased from J.T. Baker. Glycerol (cat: 65516-500ml), protocatechuate 3,4-dioxygenase pseudomonas (PCD) (cat: P8279), 3,4-dihydroxybenzoic acid (PCA) (cat: 37580-25G-F) and (+−)-6-hydroxy-2,5,7,8-tetra-methylchromane-2-carboxylic acid (Trolox) (cat: 238813-5 G) were ordered from Sigma. 1× Phosphate Buffered Saline (PBS) pH 7.2 (cat: 10010-023) was purchased from Gibco. Bovine serum albumin (cat: 001-000-162) was ordered from Jackson ImmunoResearch. Triton X-100 (cat: T8787-50ML) was purchased from Sigma Aldrich.

### DNA plasmids

A plasmid containing codon-optimized cDNA encoding human VPS13B, also including mScarlet fluorescent protein after amino acid residue 1301 flanked by BamHI restriction enzyme sites, was generated by and purchased from GenScript Biotech. This plasmid was linearized with BamHI and used to clone VPS13B^EGFP and VPS13B^Halo by In-Fusion Cloning (Takara Bio). FAM177A1-GFP, FAM177A1-RFP, FAM177A1-Halo and FAM177A1-SNAP are all cloned fromFAM177A1 pcDNA3.1+/C-(K)-DYK (GenScript Clone ID:OHu30351D) using the primers in (Table S1) by HiFi (NEB) cloning. ZFPL1-GFP was purchased from GenScript (Clone ID: OHu29409C, Accession No.: NM_006782.3). GalT-RFP was previously cloned from Addgene plasmid # 11929 (Dong et al., 2016).

### Cell Culture and transfection

Cells were cultured at 37 °C and 5% CO2 in Dulbecco’s Modified Eagle Medium (DMEM) containing 10% Fetal Bovine Serum, 100 U/mL penicillin, 100 mg/mL streptomycin, 2 mM L-glutamine (all from Gibco) and Plasmocin prophylactic 5 µg/mL (Invivogen). COS7 and HeLaM cells for imaging experiments were seeded on glass-bottomed dishes (MatTek) at a concentration of 50 to 75 × 103 cells per dish, transiently transfected using FuGene HD (Promega), and imaged ∼48 h later.

A detailed description of cell culture, transfection, immunocytochemistry, and imaging can be found in protocols.io dx.doi.org/10.17504/protocols.io.eq2lyp55mlx9/v1

### Microscopy

#### Live cell imaging

Just before imaging, the growth medium was removed and replaced with prewarmed Live Cell Imaging solution (Life Technologies). Imaging was carried out at 37 °C and 5% CO2. Spinning-disk confocal microscopy was performed using an Andor Dragonfly system equipped with a PlanApo objective (63×, 1.4 numerical aperture, oil) and a Zyla sCMOS camera, airyscan imaging is performed using Zeiss LSM 880 Airyscan. Images were analyzed in FIJI.

#### FLASH-PAINT

##### Imaging Buffer

Buffer C (1xPBS, 500mM NaCl). The imaging buffer was supplemented with 1× Trolox, 1× PCA and 1× PCD (see paragraph below for details).

##### Preparation of Trolox, PCA and PCD

100× Trolox: 100 mg Trolox, 430 μL 100 % Methanol, 345 μL 1M NaOH in 3.2 mL H_2_O.

40× PCA: 154 mg PCA, 10 mL water and NaOH were mixed and pH was adjusted 9.0.

100× PCD: 9.3 mg PCD, 13.3 mL of buffer (100 mM Tris-HCl pH 8, 50 mM KCl, 1 mM EDTA, 50 % Glycerol). All three were frozen and stored at -20 °C.

##### Cell fixation preserving Golgi complex

Cells were fixed with 4% PFA for 30 min. After four washes (30 s, 60 s, 2× 5 min), cells were blocked and permeabilized with 3% BSA and 0.25% Triton X-100 at room temperature for 1 h. Next, cells were incubated with the anti-GM130 antibody, anti-COPI antibody, and the GFP-nanobody in 3% BSA and 0.1% Triton X-100 at 4 °C overnight. Additionally, all other primary antibodies were pre incubated with the corresponding nanobodies (Tables S2) at 4 °C overnight. The next day, after four washes (30 s, 60 s, 2× 5 min) cells were incubated with the secondary nanobodies corresponding to GM130 antibody and COPI antibody for ∼2 h at room temperature. Next, unlabeled excess secondary nanobodies (to block unlabeled epitopes) were added to pre-incubation antibody-nanobody mixes at room temperature for 5 min. Next, the antibody-nanobody solutions were pooled and the cells were incubated with the pooled antibody-nanobody mix for ∼2.5 h at room temperature. After four washes (30 s, 60 s, 2× 5 min), the sample was post-fixed with 3% PFA and 0.1% GA for 10 min. Finally, the sample was washed three times with 1× PBS for 5 min each before adding the imaging solution.

##### Super-resolution microscope

Fluorescence imaging was carried out on an inverted Nikon Eclipse Ti2 microscope (Nikon Instruments) with the Perfect Focus System, attached to an Andor Dragonfly unit. The Dragonfly was used in the BTIRF mode, applying an objective-type TIRF configuration with an oil-immersion objective (Nikon Instruments, Apo SR TIRF 60×, NA 1.49, Oil). As excitation laser, a 561 nm (1W nominal) was used. The beam was coupled into a multimode fiber going through the Andor Borealis unit reshaping the beam from a gaussian profile to a homogenous flat top. As dichroic mirror a CR-DFLY-DMQD-01 was used. Fluorescence light was spectrally filtered with an emission filter (TR-DFLY-F600-050) and imaged on a scientific complementary metal oxide semiconductor (sCMOS) camera (Sona 4BV6X, Andor Technologies) without further magnification, resulting in an effective pixel size of 108 nm.

##### Imaging conditions

Imaging was carried out using the corresponding Imager (Table S3, S6), Adapter (Table S4, S6) and Eraser (Table S5) in imaging buffer. 30,000 frames were acquired at 25 ms exposure time. The readout bandwidth was set to 540 MHz. Laser power (@561 nm) was set to 80 mW (measured before the back focal plane of the objective), corresponding to ∼1.8 kW/cm^2^ at the sample plane. After imaging, the sample was subsequently washed three times with 200 µL of 1× PBS (on the microscope) followed by an incubation with the corresponding eraser (at 20 nM) for 3 min. This process was repeated iteratively for seven sequential rounds.

#### Image processing and analysis

DNA-PAINT data was reconstructed, postprocessed (drift correction and alignment of imaging rounds) and rendered with the Picasso package (https://github.com/jungmannlab/picasso.git) (Schnitzbauer et al., 2017).

### Fractionation of FAM177A1-GFP expressing cells

HeLa cells are cultured and transfected as described above. 24hrs after transfection, cells are washed three times with ice cold PBS following lysis with ice cold RIPA buffer. Lysates were ultracentrifuged in a benchtop ultra-centrifuge at 50,000 K for 1hr at 4 °C, after the centrifugation the supernatant and the pellet were separated. Samples were solubilized in 4X Laemni buffer for western blot analysis.

### Generation of VPS13B and FAM177A1 KO HeLa cells

sgRNAs targeting the human VPS13B (gRNA1 5’-CAAAATCATCAATCAAACCG and gRNA2 5-AGTGAAAGCTGTAGATCCGA) and FAM177A1 (guide RNA 5’-ATATAGATGAGTAACGAAAG) genes were generated using the IDT-DNA and Synthego online tool. HeLa cells were transiently transfected using FuGene HD (Promega) with plasmids containing Cas9 and the sgRNAs [the backbone used was plasmid PX458, a gift from F. Zhang (Broad Institute, Cambridge, MA; Addgene #48138)]. Transfected GFP expressing cells were then selected by FACS after 48 hrs via GFP expression and clonal cell populations were isolated approximately a week after.

Mutations in the FAM177A1 and VPS13B gene were confirmed by PCR and sequencing using the primers listed in (Supplemental table 1) and by western blot. A detailed description can be found in protocols.io dx.doi.org/10.17504/protocols.io.5jyl85x89l2w/v1

#### BFA treatment

70-80% confluent HeLa cells were treated with 5 µg/mL final Brefeldin A (Sigma Aldrich Catalog no # B7651) for 1hr at 37 °C. BFA was washed out by three brief rinses in PBS and cells were then incubated at 37 °C for indicated times before fixation with 4% PFA (EMS, catalog no: 15710). For the quantification of Golgi complex recovery following BFA treatment the Ilastik 1.4.OS machine learning software was used. The software was trained to recognize compact perinuclear GM130 signal (assembled Golgi complex) versus disperse GM130 signal (disassembled Golgi) using an independent training dataset. Maximum intensity projections of unprocessed images of the GM130 signal were used. A minimum of 75 cells per condition are counted. **Image processing, analysis, and statistics.** Statistical analysis was performed with GraphPad Prism 7 software. Groups were compared using a two-tail unpaired Student t-test, and results were deemed significant when a P value was smaller than 0.05.

### Zebrafish husbandry

All zebrafish studies reported here were approved by the Institutional Animal Care and Use Committee at Washington University in St. Louis. Zebrafish at all developmental stages were maintained at 28.5 °C unless otherwise specified using the standard operating procedures and guidelines established by the Washington University Zebrafish Facility, described in detail at https://zebrafishfacility.wustl.edu/facility-documents/. AB* and *fam177a1a^stl700/stl700^*; *fam177a1b^stl746/stl746^* (*fam177a1a/b DKO*) (Legro et al., in revision) zebrafish lines were used in this study.

### Annotation of zebrafish vps13b

RNA-seq reads from zebrafish Nadia strain ovary and testis (PRJNA504448) were aligned to the Nadia genome assembly with STAR (Dobin et al., 2013). Sequences were manually annotated from the alignments using Apollo v 2.4.1. (Dunn et al., 2019).

### Quantitative RT-PCR

Each RNA sample was isolated using RNeasy Micro Kit (QIAGEN, #74004) from 30 wild-type (WT) embryos. cDNA was synthesized with the iScript kit (Bio-Rad, #1708841) using 1mg of total RNA. qRT-PCR reactions were set up using SsoAdvanced SYBR green (Bio-Rad, #1725270) on CFX Opus 96 (Bio-Rad) and the gene expression was analyzed applying CFX Maestro program (Bio-Rad). Primers used are listed in Supplementary materials Table S1.

### Generation of *vps13b* crispants

#### gRNA design

To generate *vps13b* crispants, two guide RNAs (gRNAs) targeting exon 23 of the zebrafish *vps13b* gene were designed using the CHOPCHOP tool (http://chopchop.cbu.uib.no (Montague et al., 2014)). The target sequences of the gRNAs are 5’-ACTGCCGTCTCCCAGTACGC and 5’-TCCCGGGCACGGTGCGCAGC. The gRNAs, comprising a duplex made of a target sequence-specific crRNA (CRISPR RNA) and a tracrRNA (trans-activating CRISPR RNA), were prepared following the manufacturer’s protocol (IDT). Ribonucleoprotein (RNP) mix was assembled with HiFi Cas9 (IDT, #1081060) and approximately 33pg of each gRNA and 5ng of HiFi Cas9 were co-injected into the early one-celled WT and *fam177a1DKO* zygotes. At 25 hours post fertilization (hpf), we analyzed overall morphology of the injected embryos, and a randomly selected subset of these embryos was assayed for the target locus disruption through genotyping PCR using *vps13b*_geno_F/R primers (see Supplementary materials Table S1) and resolving the resulting PCR products on Fragment Analyzer (Agilent).

#### BFA treatment

BFA (Sigma-Aldrich, #B7651) was dissolved in DMSO and diluted in egg water (60 mg/mL Instant Ocean Sea Salt in distilled water) at 0.8mg/ mL. Zebrafish embryos at 6 hpf were incubated in BFA solution until 25 hpf.

## Supplemental Figure Legends

**Supplemental Figure 1.**
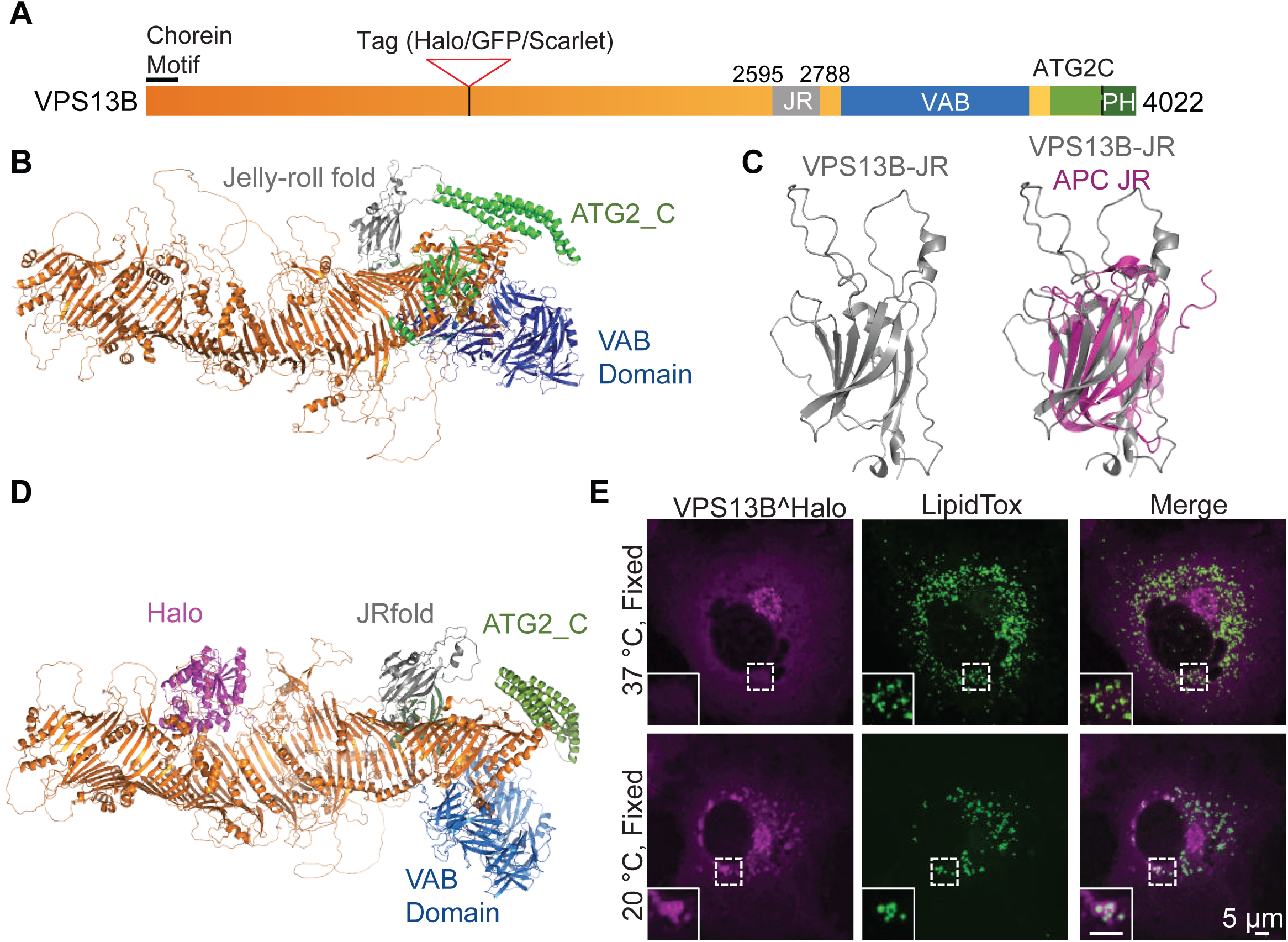
Structure and localization of VPS13B. **(A)** Domain structure of human VPS13B. **(B)** Alphafold2 predicted structure of full length human VPS13B, orange indicates the rod consisting of 13 RBG domains. **(C)** Alphafold2 prediction of the Jelly Roll domain of VPS13B alone on the left and overlayed with the Jelly Roll Fold of anaphase promoting complex subunit Doc1p/Apc10 on the right. **(D)** Alphafold2 prediction of VPS13B internally tagged with Halo tag (magenta). **(E)** COS7 cells expressing VPS13B^Halo and kept at 37°C (top panel) or shifted to 20°C for 30 minutes before fixation (bottom panel) and then stained with Lipid Tox. Scale bar = 5 µm.

**Supplemental Figure 2.**
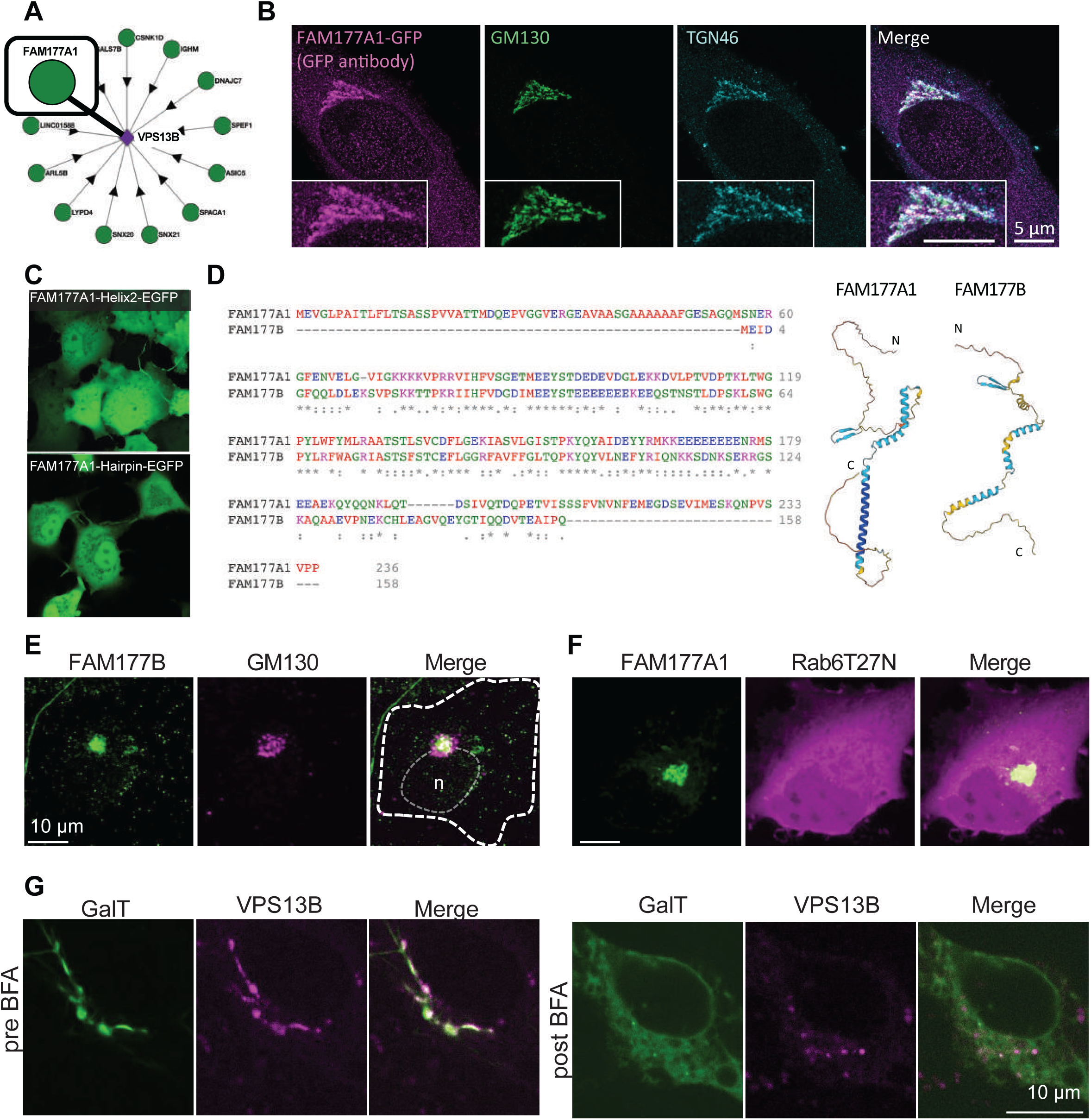
Human FAM177A1 and FAM177B localize at the Golgi complex. **(A)** Bioplex data showing predicted partners of VPS13B. (https://bioplex.hms.harvard.edu/explorer/network.php). **(B)** HeLa cells expressing FAM177A1-GFP immunolabeled with anti-GFP, anti-GM130 and anti-TGN46 antibodies. **(C)** COS7 cells expressing FAM177A1-helix2-GFP (top) or FAM177A1-Hairpin-GFP (bottom). **(D)** Left panel, Sequence alignment of FAM177A1 and FAM177B. Right panel, Alphafold2 predicted structure of FAM177A1 and FAM177B. **(E)** HeLa cells expressing FAM177B-flag fixed and immunolabeled with anti-flag and anti-GM130 antibodies. Scale bar = 10 µm. **(F)** HeLa cells expressing FAM177A1-GFP and Rab6T27N-RFP. Scale bar = 10 µm. **(G)** HeLa cells expressing GalT-RFP and VPS13B^Halo before (left panel) and after BFA treatment (5 µg/mL for 40 minutes, right panel). Scale bar = 10 µm.

**Supplemental Figure 3.**
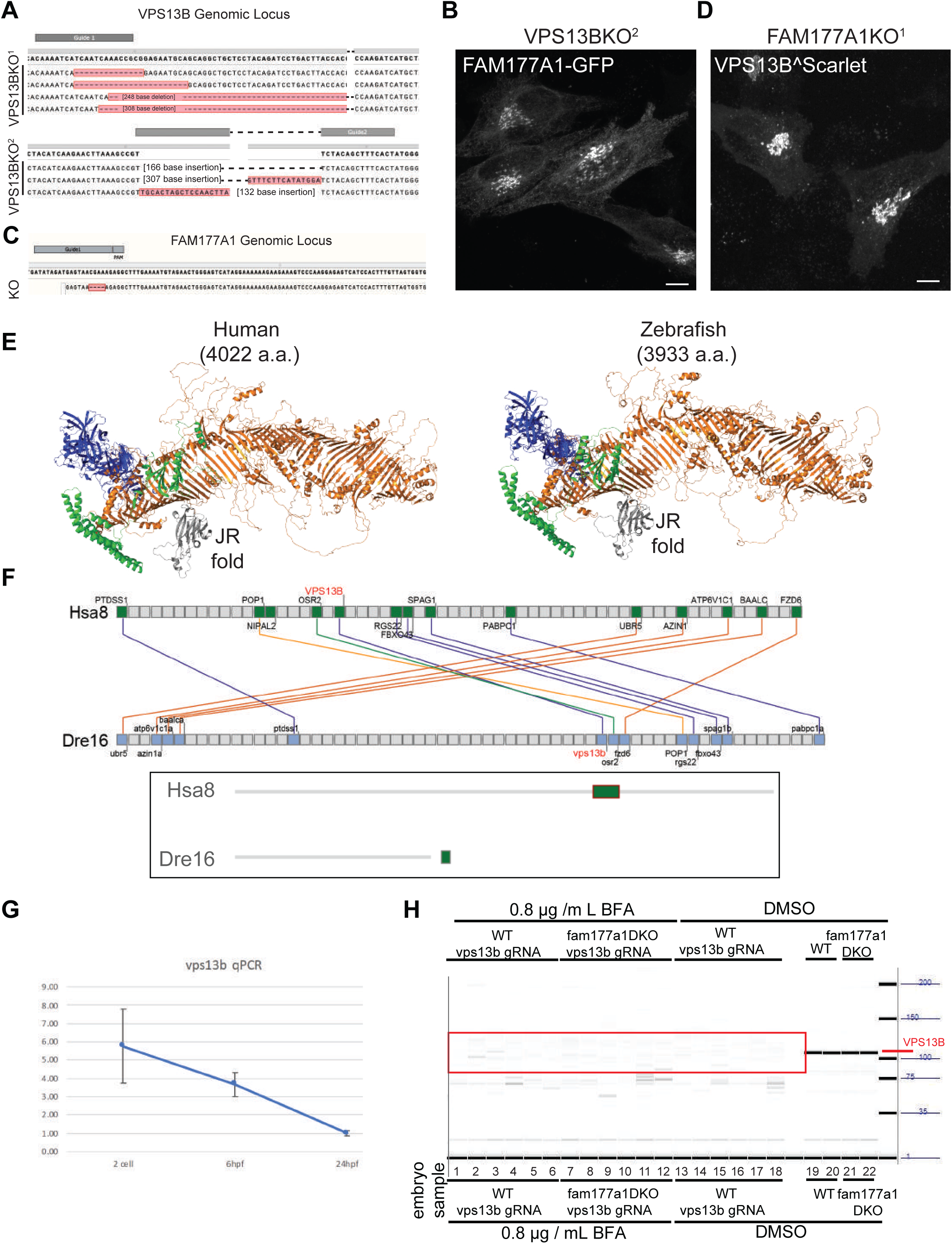
Generation of VPS13B and FAM177A1 KO HeLa cells and Zebrafish genome encodes and expresses a VPS13B homolog. **(A)** Sanger sequencing of *VPS13BKO^1^* and *VPS13BKO^2^*HeLa cells; superscripts indicate different clones. **(B)** *VPS13BKO^2^*HeLa cells expressing FAM177A1-GFP **(C)** Sanger sequencing of *FAM177A1KO* homozygous HeLa cells **(D)** *FAM177A1KO* cells overexpressing VPS13B^Scarlet. Scale bar = 10 µm. **(E)** Side-by-side comparison of Alphafold2 predicted structrure of zebrafish Vps13b and human VPS13B. **(F)** Conserved syntenies of *VPS13B* in human and zebrafish validate orthology implied by sequence comparisons. A small part of human (*Homo sapiens*) chromosome 8 (Hsa8, green part in insert) has conserved synteny with a short portion of zebrafish (*Danio rerio*) chromosome16 near its right tip (Dre16, green portion in insert). **(G)** qRT PCR analysis of WT *vps13b* in early zebrafish embryos at 2 cell stage, 6 hpf, 24 hpf. **(H)** Genotyping of *vps13b* CRISPR target locus in WT zebrafish embryos injected with *vps13b* gRNAs/Cas9 and treated with 0.8 µg/mL BFA (samples 1-6), *fam177a1a;fam177a1b DKO* embryos injected with *vps13b* gRNAs/Cas9 and treated with 0.8 µg/mL BFA (samples 7-12), WT zebrafish embryos injected with *vps13b* gRNAs/Cas9 and treated with DMSO (samples 13-18), WT zebrafish embryos treated with DMSO (samples 19,20), *fam177a1a;fam177a1b DKO* embryos injected with *vps13b* gRNAs/Cas9 and treated with DMSO (samples 21,22).

**Video 1**

Live imaging of COS7 cell in Fig.1G expressing VPS13B^halo, ZFPL1-GFP (a cis Golgi marker) and GalT-RFP (a trans-Golgi marker) during hypotonic schock.

**Video 2**

Live imaging of HeLa cell in Fig. 2F expressing FAM177A1 GFP and VPSB^Halo after addition of BFA (5 µg/mL).

## Supplemental Tables

**Table S1.**
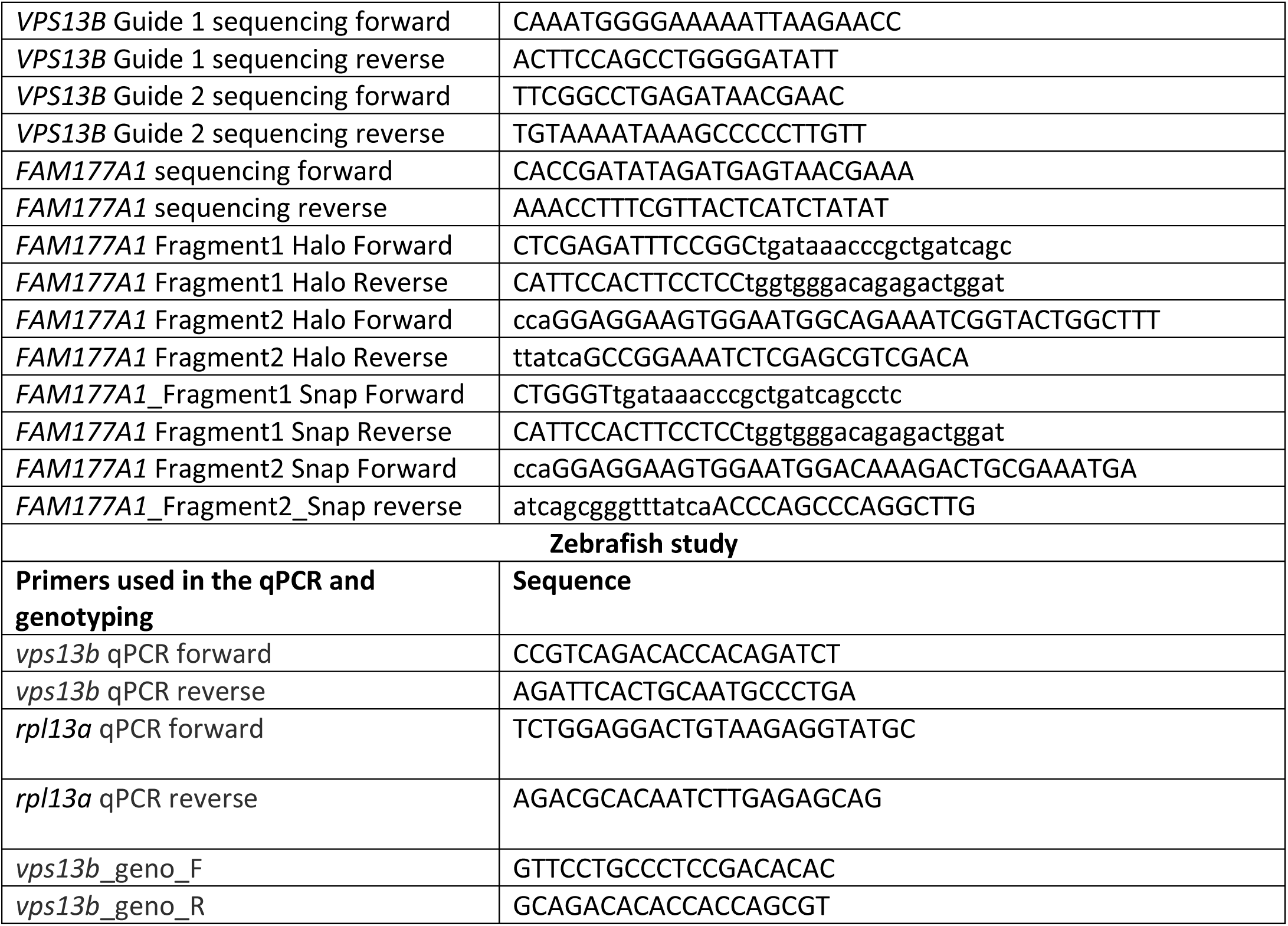
List of primers used in this study.

**Table S2.** FLASH-PAINT Nanobody Sequences.

| Target | Nanobody | Sequence (3' → 5') |
| --- | --- | --- |
| VSP13b-GFP | Anti-GFP-Nanobody-A3 | NB - TT TCTTCATTAGCG |
| TGN46 | Anti-Rabbit-Nanobody-A15 | NB - TT ATAGTGATTGGA |
| GM130 | Anti-Rabbit-Nanobody- A39 | NB - TT TTATGTTCTGCT |
| GOLGA1_1 Golgin-97 | Anti-Rabbit-Nanobody-A8 | NB - TT ATGTTAATGGGT |
| GRASP65 | Anti-Rabbit-Nanobody-A38 | NB - TT ATTTAGTGTAGC |
| GOLGB1 Giantin | Anti-Rabbit-Nanobody-A20 | NB - TT ATATGATCTCCG |
| COPI (CMIA10) | Anti-Mouse-Nanobody-A27 | NB - TT AAAAAGTTCGAG |

**Table S3.**
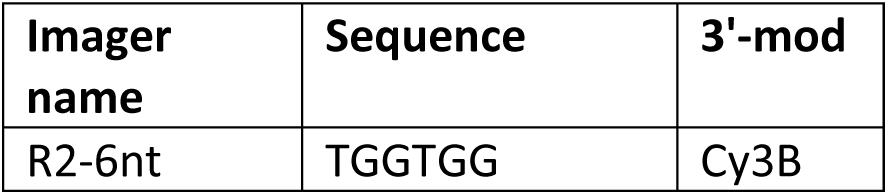
FLASH-PAINT Imager Sequence.

| Imager name | Sequence | 3'-mod |
| --- | --- | --- |
| R2-6nt | TGGTGG | Cy3B |

**Table S4.**
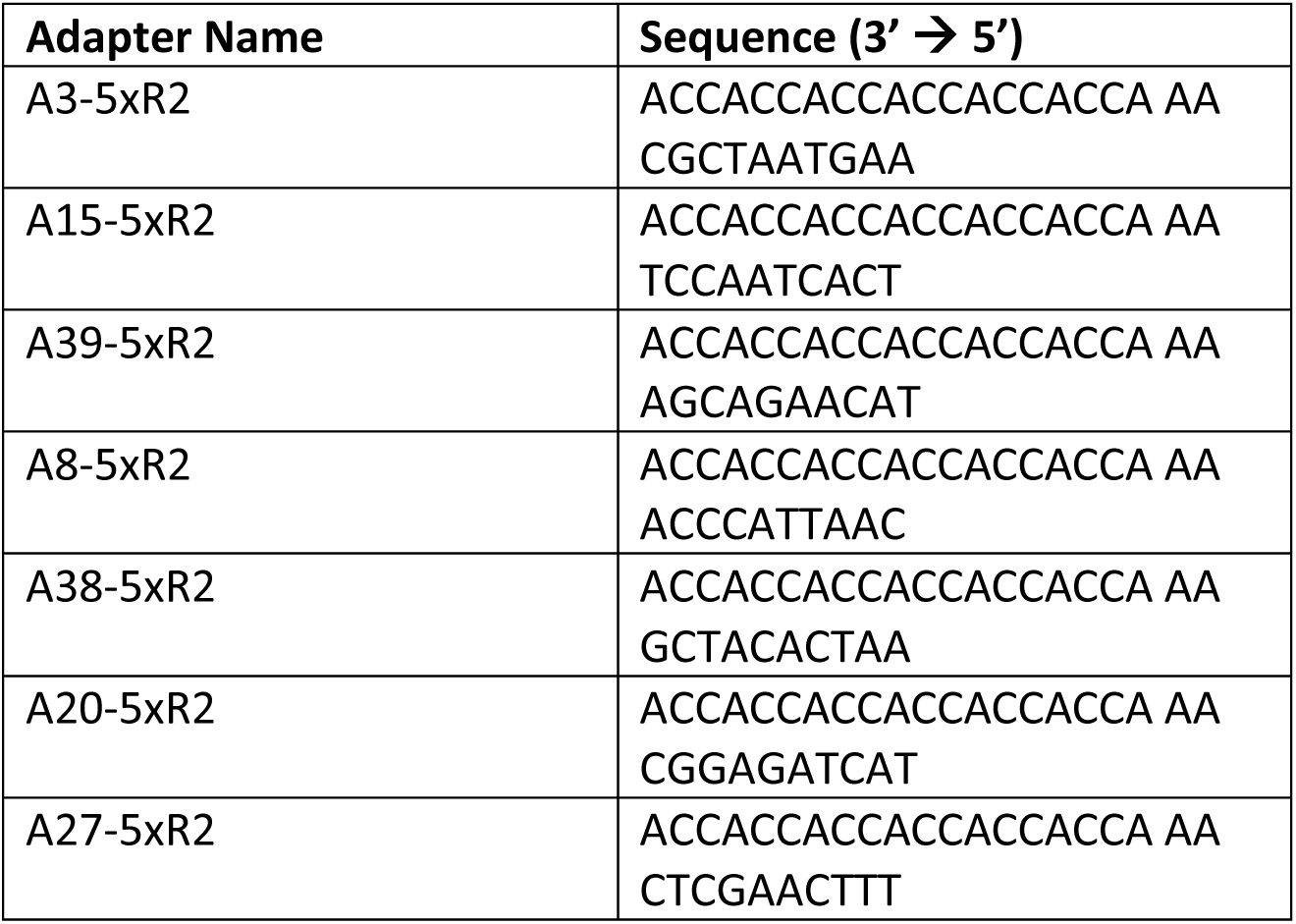
FLASH-PAINT Adapter Sequences.

**Table S5.** FLASH-PAINT Eraser Sequences.

| Eraser Name | Sequence (3' → 5') |
| --- | --- |
| E3-5xR2 | TTCATTAGCG TT TGG |
| E15-5xR2 | AGTGATTGGA TT TGG |
| E39-5xR2 | ATGTTCTGCT TT TGG |
| E8-5xR2 | GTTAATGGGT TT TGG |
| E38-5xR2 | TTAGTGTAGC TT TGG |
| E20-5xR2 | ATGATCTCCG TT TGG |
| E27-5xR2 | AAAGTTCGAG TT TGG |

**Table S6.** FLASH-PAINT Adapter and Imager Concentrations.

| # Round | Protein | Adapter –<br>Concentration | Imager –<br>Concentration |
| --- | --- | --- | --- |
| Round 1 | VPS13b | A3-5xR2 – 20 nM | R2 – 200pM |
| Round 2 | GM130 | A15-5xR2 – 20 nM | R2 – 500 pM |
| Round 3 | TGN46 | A39-5xR2 – 20 nM | R2 – 200 pM |
| Round 4 | COPI | A8-5xR2 – 20 nM | R2 – 500 pM |
| Round 5 | Golgin97 | A38-5xR2 – 20 nM | R2 – 500 pM |
| Round 6 | Giantin | A20-5xR2 – 20 nM | R2 – 500 pM |
| Round 7 | Grasp65 | A27-5xR2 – 20 nM | R2 – 300 pM |

## Supplemental Sequences

### >vsp13b predicted peptide sequence

MLESYVSPLLMSYVNRYIKNLKPSDLQLSLWGGDVVLSKLDLRLDVLEQELKLPFTFLSGHIHELRIHVPWTKLSSEPVVVTINTMECILKLRDGATDDSESCASSSTSRSAPEGSRAAVRARRQQQQQQGPSGPDLPPGYVQSLIRRVVNNVNIVVNNLILKYVEDDIVLSVNITSAECYTVDELWDRAFMDISAPELVLRKVINFSDCTVCLDRRNASGKIEFYQDPLLYKCSFRTRLHFSYDSINAKIPAVIKIHTMVESLKLSLTDQQLPMFLRLLELGVALYYGEMEERREVEREDSGPLRDSIPGLRDGQCDEDEDGDEDGEQGWVSWAWSFIPAMVSAEEEERERAEGEQGAEFVECLEPGAPRPLSTAHRDPVVSIGFYCTKASVTFKLTESCSESSFYSPQKLKSREVLSVEQEGITVEALMMGEPFFDCQVGVVGCRALCLKGIMGVRDFEDNMNRKEEDAVFFRCGDTLSLKGMTYLTNSLFDYRSPENNGVRAEFILEGNLHKETYTEAAGLQRYGAFYMDYLYMMESSSRVCAGLQDGAALAVRLQETSLKRVVFGALDLQLHSSAVHRMLKMITCTLEHEYQPYCRAQPPDVVEECVSVDPEQVAALEECIPVRQTIVTLMRATVTIPAAEYNLLHLILPTLLGHKVTSAQTSVPQFQLLRPLPALRLQFQRVTFEHSQPMHEEEVTRAASSLKQPSHTLLHHCYTHCCLKVFELQAGLTVIGSEEASQPLIPIIPAFSTALYGKQLHLPAYWLRNPSVQVSECVLELPQVCVQATRAQVLLLHCMTRSWTHSEGGGACSGITDSLISHAYNTTGVKSCAPVLEVCVQRVELKVCVRPALLCVSGTLGAVKVCARTPGVCEGQKDQLVPLLQGPSDTTDLHSSRWLSGSRKPASLLSPDLLQLTVQLPQQEHTPNPGAVLLLSVQGIAVNLDPVLSSWLLFHPQRTSGSRQTQQVSMAMKKRREDEASVGSAALTRQASNQNSDYTSSPVKTKTVTESRPLSIPVKVFPSAEECPVSPEEQMKNLITHTWNAVKHLTLQVELQSCCVFLPSDTLPSPSTLVCGDVPGTVRSWYHSQVCMPGTLVVCLPQISVLSAGHRRMEPLQDGPLTVPRPVLEEGGAFPWTVCVRQLSVYSLLGHQRSLSVVDPLGCTSTLALTAPRLQPAARDAFIICLHVDLQPLQLQCSNPQVQLVCALWCRWMQIFSVLERLQTRGAQRAAAGFPECSAPAAGPASPVHSSAGTAPPDTSTCSPSADLGTPTEADSAHTDDPAAGETLTLEQQTCSISGSSRRLSVWMQWMLPRLTLKLFSSEPASRTTELCVLAEMEDLSASVDIQDVYTKVKCKVGSFHIDHYRSSAEEAAVVLSCTEKLNRRTVLLRPLGRQEPHSAFSFFPPAAAKALEVSHQQHGFLSVTYTQAVTRNVRHKLTARQERMAATPRLSEDTSDASPQHLHEILLTAQPFDLVLSCPLLAAVARVFHLPPSLRPPTRERISAQPMRGHALSSSSLPLVYVNTSVIRIFCPLEDTHGSTHSQSKHWKEDTVVLKIGSVSVAPQADNPLPRSVLRKDIYQRALNLGVLRDPGSEVEDRQYQIDLQCMNIGTAQWAELQPEGAGPGGVTSADTERSSQNPALEWNMASSIRRQQEKRVILSPIITDFSVRITAAPAVIYRKPLSADHGPAEEVVVCGHSLEMNITSNLELFLSAAQVQLLQRLLQDNMRLTEDADTTADVCGQQQQQAAAVSGCVESVCVGGAPCGQDSGFGSDSARLRIVQHRLARPPQQPTVTKSLSFVPFDVFLTAGRISLMTYATAPTDPADPTAPASPPAPPAPAVQDGGLASLTADSLLRGGSCMSAPGRSSARQALGVTVVRQPGRRGDTHTCLQPLLLLQVMQPSVLLSCQHRRQRLELSLFDLSLKGPVADYRSHDAGKSLPESLDYSVFWLQTVAGEADGRTGIPPPLLSVSVRDFLSAPELQVEVSRPLRVSPTPAKVEQAKLFWRTLFPEGEALPTDKTGPTPSCSEAPPTVDRLLGALQTPVPFRRLALHAVQLVLSVETQTCSSVTLSVSALTSTLTLQNNSSSRPADGFKEVCVSLQCEDLLIRTALRERSSVFVGPFSCSADLEAHWCRHSGSSAPDAPGPPRVLIDMKGGLLQVFWGQLQFNCLSEFLEQLQNYWSQLSTVEAEPQDKPCPSMPPPTSLSAQSEHSSDDLRTGLFQYIQDSACQRLPAPHEVVFWRETEDSPGVMLWRYPEPRAITFLRITPVPFNTTEDPDISTADLGDVLQVPCSLEFWDELQRVFVPYREFSLSESSVCELTLPTLTPDTQQTDLVTSDLWRMVLNSNSDGGDESSDSESGSQVHCEQLVCPTALAACTRVDSCFAQWCVSSVGVSMRLAYLQLHFCHNLDQLGTVPCQKLRPFLPDRKLPQDQEFAVVCVREPCVFVRQWCGVVQSCVELSLSCALSCSLLEYRNLTLLPVLQPTRMQAHATHTLTHTHAQHTLHCHATLQPLHAAIGQYAIHTLDRALQAWRQNGVSDAEEVVFCHYVICNDTQEVLRFGQVDTDENILLQSNQSHQYSWRTHRSPQLLHICMEGWGNWRWAEPFSVDDAGTLLRTIQHRGQTASLIIRVTQLSAVQKQVVISGRQVVCSFLHQPIELRLLQRVCVDGQPLLRARVCRLEPDCRPPAFVLEHAELTEVCVRACGEDAWSQDVCLEHSDTHNSSVVQVPSSNGSLLHVWCTRVLLEPNTHTQQRVVVFSPLFMMWSHLPEPVLVHVEKRSLGLRDTQLIPGRGHQEALLNAEADLTHHLTFQAREEEGASHCAVPVSTAVVKQILSRASAEHQQNPQNILEHFCGEKKPSRAAWPYSSREAERGAGEPVAQWDSPMQVRLSVWRAGLNTLLVELLPWALLVNRSHCDLWLFEAENIIVQIPAGGTIVPPNLTDAFQIGIYWPHTNTVHKSPAVQLVHEVSSPRWPEGGGADVLLLDEEGFIHTDITLGTQPGNLKLCQFCVSSSVKFGIQVLQIEDKTVLVNNTTHTFKCRAVIPQHTAVTPAQLCPVPESSVFALGPAGAAGDQVCCALPCWDVLRGPSSEDPPLGPQLLQLSCEGCVLRWSPAAPVRSDRPRQSLPVPVEPSSDWPFSTRPLVLTCQEHLGVTYLTVSEDQSPRMLIHNHCPKSLLLKENTRECVRWPVFPRVLPAHSSVHHELLHQASSFPECRQKETLPTLRLRILTDTSTGPTDTPTHSTDTPTDPTDTPTTPGWTEAIDISSPGTQVVFLPGFGCLYIDVLQQSGSVTLTLAAESSAAELITQHRPCRQMLSFRLLLDEASVALCDDITSPSGSVELLRLTISKLLLLLPPCEPPADPSTAPALHTLQVLCGGLQVDNQLYERASFHFPVMVCQEPQGEAELCPQQLQQFCAACFLSLSISVCADGQLDRLALRIRPARLYLEDTFIYYMKTLFHTYIPECAAGGRGCVRGVESAVPQEVLESMRALVSPLRLQKLSIEPLQLLVSIHASLKLYIASDHTPLAFSEFERGPVCTTARHLVHTLAMHYAAGALFRAGWVVGSLEILGSPASLVRSIGNGVSDFFRLPYEGLTRGPGAFISGVSRGTNSFIKHISKGTLTSITNLATSLARNMDRLSLDDEHYTRQEEWRRQLPETLGDGLRQGLSRLGISLLGAVAGIVDQPMQTFTRTLELPNSASSAARGVISGVGKGIVGVFTKPIGGAAELVSQTGYGLLHGAGLWQLPKQLHQPTDNRSADAANSSAKYIWKMLQSLGRAELHMALDVCMVSGSGQERAGCLLLSAEVLFVVSVCEDAQQQAFPITEIQCEHDTHTPGRITLTLQQQRVNTDAEVEGVRERLSELQYTRLLDFVRGASPFLPAALQPSVSPAEPQRSVCRTYQYQADAAFARVFVCRFQMVKNRALRIGFH

### >vsp13b predicted cdna sequence

TGACGCGTGGACGCAGCCTGTGTGTGCGGAAGTAATGGTCGTGGTGTGTGTTTGTGTATGTGGAGCAGCCGGACGCGTCGCATGAACCCGCGTGAGGGTGTGTCAGGATGTTGGAGTCGTATGTGTCTCCTCTGCTGATGAGTTATGTGAACCGCTACATAAAGAACCTGAAGCCGTCAGACCTGCAGCTGTCACTATGGGGCGGCGACGTGGTGCTGAGCAAACTGGACCTGCGGCTGGACGTGCTGGAGCAGGAGCTGAAGCTGCCCTTCACCTTCCTGAGCGGCCACATCCACGAGCTGCGCATCCACGTGCCCTGGACCAAGCTGAGCTCAGAGCCGGTGGTCGTCACCATCAACACCATGGAGTGCATCCTGAAGCTGCGGGACGGAGCCACGGATGACTCTGAGAGCTGTGCCTCCAGCTCCACCAGTCGCAGTGCTCCTGAGGGCTCCAGAGCTGCAGTGAGGGCCCGGCGACAGCAGCAGCAGCAGCAGGGCCCCAGCGGCCCCGACCTGCCCCCAGGCTATGTGCAGAGTCTGATCCGGCGTGTGGTCAATAATGTGAATATCGTGGTCAATAACCTGATCCTGAAGTACGTGGAGGACGACATCGTGCTGTCGGTGAACATCACGTCTGCGGAGTGTTACACGGTGGACGAGCTGTGGGACCGGGCCTTCATGGACATCAGCGCTCCAGAACTGGTCCTCCGGAAGGTGATCAACTTCTCGGACTGCACCGTGTGTTTGGACCGCCGGAACGCCAGCGGGAAGATCGAGTTTTACCAGGACCCGCTGCTCTACAAGTGCTCCTTCAGGACCCGTCTGCACTTCAGCTACGACAGCATCAACGCTAAGATCCCCGCCGTCATTAAAATTCACACTATGGTGGAGAGTCTGAAGCTCTCCCTGACGGATCAGCAGCTGCCCATGTTCCTCCGGCTGCTGGAGCTGGGTGTAGCGCTGTATTATGGGGAGATGGAGGAGCGGCGGGAGGTGGAGCGGGAAGACAGCGGCCCACTGCGAGACAGCATTCCCGGGCTGCGGGATGGGCAGTGTGATGAGGATGAGGATGGGGATGAGGATGGGGAGCAGGGCTGGGTGTCGTGGGCCTGGTCCTTCATCCCAGCCATGGTGAGCGCTGAGGAAGAGGAGCGCGAGCGTGCGGAGGGTGAGCAGGGTGCAGAGTTTGTGGAGTGTTTGGAGCCCGGAGCCCCGCGGCCCCTCAGCACAGCACACAGAGACCCCGTAGTGTCCATCGGCTTCTACTGCACTAAAGCCTCCGTCACCTTCAAGCTGACGGAGAGCTGTTCTGAGAGCAGTTTCTACAGTCCGCAGAAGCTGAAGTCTCGAGAGGTGCTGAGCGTGGAGCAGGAGGGCATCACTGTGGAGGCGCTGATGATGGGCGAGCCGTTCTTCGACTGTCAGGTGGGCGTGGTGGGCTGCAGAGCGCTGTGTCTGAAGGGCATCATGGGTGTGCGAGACTTTGAAGACAACATGAACAGAAAGGAGGAGGATGCTGTGTTCTTCCGCTGCGGTGACACACTGAGTCTGAAGGGCATGACGTATCTGACCAACTCACTGTTCGACTATCGGAGTCCAGAGAACAACGGAGTCCGCGCAGAGTTCATCCTGGAGGGAAACCTGCACAAGGAGACGTACACAGAGGCTGCAGGTCTGCAGAGATACGGAGCGTTCTACATGGATTACCTGTACATGATGGAGAGCAGCAGCAGAGTGTGTGCGGGTCTGCAGGATGGCGCAGCTCTGGCGGTGCGGCTGCAGGAGACGTCGCTCAAGCGGGTTGTGTTCGGGGCTCTGGATCTGCAGCTCCACAGCAGCGCCGTACACCGCATGCTGAAGATGATCACATGCACACTGGAGCACGAGTACCAGCCCTACTGCAGAGCGCAGCCGCCCGATGTGGTGGAGGAGTGTGTGTCTGTGGACCCTGAGCAGGTTGCAGCGCTGGAGGAGTGTATTCCCGTGCGTCAGACCATCGTGACTCTGATGAGGGCGACGGTCACCATACCTGCAGCCGAGTACAACCTGCTGCACCTCATCCTGCCCACACTACTGGGACACAAGGTCACCTCTGCGCAGACGTCAGTCCCTCAGTTCCAGCTGCTGCGTCCTCTGCCTGCGCTGCGGCTGCAGTTCCAGCGCGTCACGTTCGAGCACTCGCAGCCGATGCACGAGGAGGAAGTGACACGCGCAGCCAGCAGCCTCAAACAGCCCTCACACACACTGCTGCACCACTGCTACACACACTGCTGCCTTAAGGTCTTTGAGCTCCAGGCGGGTCTGACTGTGATTGGCTCAGAGGAGGCGTCGCAGCCGCTCATACCCATAATTCCTGCTTTCAGTACTGCTCTCTATGGGAAGCAGCTCCACCTGCCTGCGTACTGGCTCAGGAATCCGTCAGTGCAGGTCTCCGAGTGTGTGTTGGAGCTGCCGCAGGTGTGTGTTCAGGCCACGCGCGCGCAGGTGCTGCTGCTGCACTGCATGACACGCAGCTGGACACACAGTGAGGGGGGCGGAGCCTGCAGCGGCATCACTGACAGTCTCATTAGCCACGCCTACAACACTACAGGTGTGAAGTCGTGCGCTCCTGTGCTGGAGGTGTGTGTTCAGCGTGTGGAGCTGAAGGTGTGTGTGCGGCCGGCGCTGCTGTGTGTGTCTGGAACTCTGGGAGCCGTGAAAGTGTGTGCCAGAACGCCCGGTGTGTGTGAAGGGCAGAAGGATCAGCTGGTGCCGCTGCTCCAGGGTCCGTCAGACACCACAGATCTGCACAGCAGCCGCTGGTTGAGCGGGAGCCGTAAGCCTGCGTCTCTGCTGTCCCCCGACCTCCTGCAGCTCACTGTGCAGCTGCCGCAGCAGGAACACACACCCAACCCCGGTGCTGTTCTGTTGCTCAGTGTTCAGGGCATTGCAGTGAATCTGGATCCGGTTCTGTCGTCCTGGCTGCTGTTCCACCCGCAGAGAACCAGCGGCAGCAGACAGACACAGCAGGTCTCCATGGCGATGAAGAAGAGGAGGGAGGATGAAGCTTCAGTCGGCAGCGCTGCACTGACCAGACAAGCCAGCAACCAGAACTCCGACTACACCAGCAGCCCAGTCAAGACCAAAACTGTGACAGAGTCCAGGCCTCTCTCTATCCCAGTAAAGGTTTTTCCCTCGGCTGAGGAATGTCCCGTGAGTCCAGAGGAGCAGATGAAGAACCTGATCACACACACCTGGAACGCCGTCAAACACCTCACACTACAGGTGGAGCTGCAGTCCTGCTGTGTGTTCCTGCCCTCCGACACACTGCCGTCTCCCAGTACGCTGGTGTGTGGGGACGTCCCGGGCACGGTGCGCAGCTGGTACCACAGTCAGGTGTGTATGCCGGGCACGCTGGTGGTGTGTCTGCCGCAGATCAGTGTGTTGAGCGCAGGACACCGCCGCATGGAGCCGCTGCAGGACGGCCCGCTCACTGTGCCCAGACCCGTGCTGGAGGAGGGCGGTGCGTTCCCCTGGACGGTGTGTGTGCGTCAGCTGAGTGTGTATTCTCTGCTGGGTCATCAGCGCTCTCTCAGTGTTGTGGATCCTCTGGGCTGCACCTCCACACTGGCCCTCACTGCACCCCGACTGCAGCCGGCCGCCCGAGACGCCTTCATCATCTGCCTGCATGTGGACCTGCAGCCGCTGCAGCTGCAGTGCTCCAACCCACAGGTCCAGCTGGTGTGTGCTCTGTGGTGCCGCTGGATGCAGATCTTCAGTGTGTTGGAGCGTCTGCAGACTCGAGGAGCTCAGAGAGCGGCTGCGGGTTTCCCAGAATGCTCTGCTCCAGCCGCTGGCCCCGCCTCCCCTGTGCACAGCAGCGCCGGCACCGCCCCGCCCGACACCAGCACCTGCAGCCCGTCTGCTGACCTGGGCACGCCCACTGAGGCTGACTCTGCGCACACAGACGACCCCGCGGCCGGCGAGACCCTGACCCTGGAGCAGCAGACCTGCAGCATCAGCGGCTCCAGCAGGAGACTCAGCGTCTGGATGCAGTGGATGCTGCCGAGACTCACACTCAAGCTGTTCTCCAGCGAGCCCGCGAGCAGAACCACCGAGCTGTGTGTGCTGGCAGAGATGGAGGACCTGAGCGCGTCTGTGGACATTCAGGACGTTTATACCAAGGTGAAGTGCAAAGTGGGAAGCTTCCATATAGATCACTACAGAAGCAGTGCTGAAGAGGCTGCGGTCGTCCTCTCTTGCACTGAAAAGCTAAACAGACGCACGGTTCTGCTGCGGCCGCTCGGCAGACAGGAGCCGCACAGCGCTTTCAGCTTCTTTCCCCCTGCGGCAGCGAAGGCTCTGGAGGTCTCGCACCAGCAGCACGGCTTCCTGTCGGTGACCTACACGCAGGCAGTGACGCGGAACGTGCGGCACAAGCTGACGGCACGACAGGAGCGCATGGCGGCGACCCCCAGGCTGAGCGAGGACACGAGCGACGCTTCACCACAGCACCTGCACGAGATCCTGCTGACGGCCCAGCCCTTCGACCTGGTGCTGTCCTGCCCTCTGCTGGCGGCAGTGGCACGCGTCTTCCACCTGCCCCCATCACTGCGCCCACCCACCCGCGAGCGCATCTCTGCTCAGCCAATGAGAGGACACGCACTGTCTTCCAGCAGCCTGCCGCTGGTCTACGTCAACACCAGTGTGATCCGCATATTCTGCCCCCTGGAGGACACACACGGCAGCACACATTCACAGTCCAAGCACTGGAAGGAGGACACAGTTGTGCTGAAGATTGGATCAGTGAGCGTCGCTCCTCAGGCTGATAATCCACTGCCGCGCTCCGTCCTGCGCAAAGACATCTACCAACGGGCGCTGAATCTGGGCGTGCTGCGTGACCCGGGCTCTGAGGTGGAGGACCGCCAGTATCAGATCGACCTGCAGTGCATGAACATAGGGACGGCGCAGTGGGCGGAGCTACAGCCAGAGGGGGCAGGGCCAGGCGGTGTCACATCGGCGGATACAGAGAGGAGCTCACAGAACCCTGCACTGGAGTGGAACATGGCCAGCAGTATCCGCCGGCAGCAGGAGAAGCGAGTGATCCTCTCTCCGATCATCACAGATTTCTCTGTGCGCATCACAGCAGCTCCGGCCGTCATCTACCGCAAGCCCCTCTCAGCAGATCACGGCCCAGCGGAGGAGGTGGTGGTGTGTGGTCACAGTCTGGAGATGAACATCACGTCTAACCTGGAGCTGTTCCTGAGCGCAGCACAAGTGCAGTTACTGCAGCGCCTCCTGCAGGACAACATGAGGCTGACGGAGGACGCAGACACCACTGCAGACGTGTGTGGTCAGCAGCAGCAGCAGGCCGCAGCAGTGTCGGGCTGTGTGGAGAGTGTGTGTGTCGGTGGCGCCCCCTGTGGGCAGGACAGTGGTTTCGGCAGTGACAGTGCCCGCCTGCGTATCGTCCAGCACCGGCTCGCGCGCCCGCCACAGCAGCCCACCGTCACTAAGAGCCTGAGCTTCGTCCCGTTCGATGTGTTCCTGACGGCCGGCCGCATCTCCCTGATGACCTACGCCACGGCACCCACCGACCCCGCTGACCCCACCGCGCCCGCTAGCCCCCCCGCTCCCCCCGCACCGGCAGTACAGGATGGTGGTCTGGCCTCTCTGACGGCTGACAGTCTGCTGCGCGGTGGCTCCTGCATGTCAGCTCCGGGCCGCAGCTCTGCGCGTCAGGCTCTGGGGGTGACGGTGGTGCGGCAGCCGGGCCGGCGGGGGGACACACACACCTGTCTGCAGCCACTGCTGCTCCTGCAGGTGATGCAGCCATCCGTTCTGCTGAGCTGCCAACACCGCCGGCAGCGGCTGGAGCTCTCACTGTTCGACCTCTCGCTAAAGGGGCCCGTCGCTGACTACAGGAGCCACGATGCAGGTAAGTCTCTGCCGGAGTCGCTGGACTACAGTGTGTTCTGGCTGCAGACGGTGGCAGGTGAGGCAGACGGGCGCACCGGGATCCCTCCGCCACTGCTGTCTGTGTCCGTCAGAGACTTCCTGAGCGCACCAGAGCTGCAGGTGGAGGTCAGCCGGCCGCTGCGGGTCAGTCCCACACCTGCGAAGGTGGAGCAGGCCAAACTCTTCTGGAGGACACTGTTCCCTGAGGGGGAAGCCCTGCCCACAGACAAAACAGGCCCCACCCCCAGCTGCAGTGAAGCTCCGCCCACAGTGGACCGGCTGCTGGGTGCGCTGCAGACACCTGTGCCGTTCCGCAGGCTGGCACTGCATGCAGTGCAGCTGGTGCTGAGTGTGGAGACGCAGACGTGCAGCAGTGTGACGCTCTCAGTGTCTGCCCTGACCAGCACACTCACACTGCAGAACAACAGCAGCAGCAGACCCGCCGACGGGTTTAAGGAGGTGTGTGTGTCGCTGCAGTGTGAGGACCTGCTGATCCGCACGGCTCTGAGGGAGCGCAGCTCAGTGTTTGTGGGTCCGTTTTCCTGCAGTGCGGATCTAGAAGCTCATTGGTGCAGACACAGTGGAAGCTCCGCCCCTGATGCACCGGGACCGCCCAGAGTGTTGATTGACATGAAGGGGGGCCTGCTGCAGGTGTTTTGGGGTCAGCTGCAGTTCAACTGTTTGTCTGAGTTTCTGGAGCAGCTGCAGAACTACTGGAGTCAACTGAGCACTGTGGAGGCGGAGCCACAGGATAAGCCCTGCCCCTCAATGCCTCCACCCACCTCCCTCTCTGCCCAATCAGAACACTCGTCTGATGACCTGCGCACTGGGCTCTTCCAGTACATACAGGATTCAGCGTGCCAGCGGCTGCCAGCTCCTCATGAGGTGGTCTTCTGGAGGGAGACTGAGGATTCTCCAGGTGTGATGCTGTGGCGTTACCCTGAGCCGCGCGCCATCACCTTCCTCAGAATCACACCTGTGCCTTTCAACACCACTGAAGACCCCGACATCAGCACCGCAGACCTGGGGGACGTGCTGCAGGTCCCCTGTAGTCTGGAGTTCTGGGACGAGCTGCAGCGAGTGTTTGTGCCTTACCGAGAGTTCAGCCTATCAGAGAGCAGCGTATGTGAGCTGACCCTGCCCACTCTGACCCCCGACACACAGCAGACTGACCTGGTGACCTCTGACCTCTGGAGGATGGTGCTCAACAGCAACAGTGATGGAGGAGATGAGAGCTCTGACAGTGAGTCGGGCTCTCAGGTGCACTGTGAGCAGCTGGTGTGTCCGACCGCTCTGGCCGCGTGTACGCGTGTGGACTCGTGTTTCGCGCAGTGGTGTGTGTCATCAGTCGGGGTCTCTATGCGTCTGGCGTACCTGCAGCTGCACTTCTGCCACAACCTGGACCAGCTCGGCACAGTGCCATGTCAGAAGCTCCGCCCCTTCCTCCCGGACAGGAAGCTGCCTCAGGATCAGGAGTTTGCGGTGGTGTGTGTGCGCGAGCCGTGTGTGTTTGTGCGTCAGTGGTGTGGCGTTGTGCAGAGTTGTGTTGAGCTGAGCTTGTCGTGTGCGCTGAGCTGCAGTCTGCTGGAGTACCGCAACCTCACACTGCTGCCCGTCCTGCAGCCCACGCGCATGCAGGCCCACGCCACACACACACTCACACACACGCATGCGCAACACACACTGCACTGCCACGCCACGCTGCAGCCACTGCACGCCGCCATCGGACAGTACGCCATACACACACTGGACCGAGCGCTGCAGGCCTGGAGACAGAACGGCGTGTCGGATGCAGAGGAAGTAGTTTTCTGTCATTACGTGATCTGCAACGACACGCAGGAAGTGCTGCGCTTCGGACAGGTGGACACCGATGAGAACATCCTGCTGCAGAGCAACCAGAGCCACCAGTACAGCTGGAGGACACACCGATCCCCACAGCTGCTGCACATCTGCATGGAGGGCTGGGGTAACTGGCGCTGGGCTGAACCCTTCAGTGTGGACGATGCAGGAACACTGCTGAGAACCATCCAGCACCGAGGACAGACGGCGTCACTGATCATCAGAGTCACACAGCTGAGCGCAGTGCAGAAACAGGTGGTGATCAGTGGTCGGCAGGTGGTGTGTAGCTTCCTCCATCAGCCGATCGAGCTCCGCCTCCTGCAGCGTGTGTGTGTGGACGGTCAGCCGCTGTTGCGTGCGCGTGTGTGTCGCCTGGAGCCCGACTGCAGACCGCCGGCGTTCGTGCTGGAGCACGCGGAGCTGACGGAGGTGTGTGTGCGCGCGTGCGGAGAGGACGCCTGGTCGCAGGACGTGTGTCTGGAGCACAGCGACACACACAACAGCTCAGTGGTGCAGGTGCCGTCTTCTAATGGGTCACTGCTGCATGTGTGGTGCACACGAGTGCTGCTGGAGCCGAACACACACACACAGCAGAGAGTGGTGGTGTTCAGTCCTCTCTTCATGATGTGGAGTCACCTGCCGGAGCCGGTGCTGGTGCACGTGGAGAAGCGCAGTCTGGGCCTCAGAGACACACAGCTGATCCCCGGACGCGGACACCAGGAGGCGCTGCTGAACGCGGAGGCAGACCTCACACACCACCTCACCTTTCAGGCCAGGGAGGAGGAGGGCGCATCTCACTGCGCAGTTCCTGTGTCCACTGCTGTCGTCAAGCAGATCCTGAGCAGAGCATCAGCAGAACACCAGCAGAACCCGCAGAACATCCTGGAGCACTTCTGTGGAGAGAAGAAGCCCAGCAGAGCAGCCTGGCCATACAGCAGCCGAGAGGCAGAGCGGGGTGCGGGCGAGCCGGTGGCGCAGTGGGACAGCCCGATGCAGGTGCGTCTGAGTGTGTGGCGCGCGGGTCTGAACACACTGCTGGTGGAGCTGCTGCCGTGGGCTCTGCTGGTCAACCGCTCACACTGCGACCTGTGGCTGTTCGAGGCCGAGAACATCATCGTGCAGATCCCGGCCGGAGGAACCATCGTCCCGCCCAACCTCACGGATGCGTTCCAGATCGGCATTTACTGGCCGCACACTAACACAGTGCACAAGTCTCCTGCCGTGCAATTGGTGCATGAAGTGTCGTCACCACGGTGGCCAGAAGGGGGCGGAGCTGATGTGCTGCTGCTGGATGAGGAGGGATTCATCCACACGGACATCACACTCGGAACACAGCCAGGCAATCTCAAGCTCTGTCAGTTCTGTGTGTCGTCTTCAGTGAAGTTCGGGATTCAGGTTTTGCAGATTGAGGATAAAACTGTCCTGGTGAACAACACGACACACACCTTCAAGTGTAGAGCTGTGATTCCACAACACACAGCGGTCACACCTGCGCAGCTCTGTCCTGTCCCAGAATCCTCAGTGTTTGCTCTGGGCCCCGCGGGTGCGGCTGGAGATCAGGTGTGCTGTGCTCTGCCCTGCTGGGATGTGCTGCGGGGTCCGTCCTCAGAAGACCCTCCGCTGGGGCCGCAGCTCCTGCAGTTGAGCTGTGAGGGGTGTGTGCTGCGGTGGAGCCCTGCGGCCCCGGTGCGCTCCGATCGGCCACGACAGAGTCTTCCGGTGCCCGTAGAGCCCAGCTCCGACTGGCCCTTCAGCACCAGGCCTCTGGTTCTGACCTGTCAGGAGCATCTGGGCGTCACGTATCTGACAGTGAGTGAAGATCAGAGTCCACGCATGCTGATCCACAACCACTGCCCCAAGAGCCTCTTACTGAAGGAGAACACCAGAGAGTGTGTGCGGTGGCCGGTGTTCCCCCGTGTCCTCCCTGCGCACTCCTCCGTCCACCATGAGCTGCTCCATCAGGCCTCCAGCTTCCCTGAGTGCCGACAGAAGGAGACGCTTCCCACCCTGCGGCTGCGCATCCTCACAGACACGTCCACAGGCCCCACAGACACACCCACACATTCCACAGACACGCCCACAGACCCCACAGACACGCCCACTACGCCAGGGTGGACAGAAGCCATTGACATCAGCAGTCCTGGAACACAGGTGGTTTTCCTCCCTGGTTTCGGCTGTTTGTACATTGATGTTCTTCAGCAGAGCGGCAGCGTCACCCTGACACTGGCAGCAGAGAGCAGCGCAGCAGAGCTCATCACACAGCACAGGCCGTGCCGTCAGATGTTGTCCTTCCGGCTGCTGCTGGATGAGGCCAGTGTTGCGCTCTGTGATGACATCACCAGCCCGTCCGGCTCCGTGGAGCTGCTGCGCCTCACCATCTCCAAACTGCTGCTGCTGCTGCCCCCCTGTGAACCGCCCGCTGACCCTAGCACTGCACCTGCCCTCCACACGCTGCAGGTGCTCTGCGGCGGTCTGCAGGTGGACAACCAGCTGTATGAGCGTGCCAGCTTTCACTTCCCGGTCATGGTGTGCCAGGAGCCGCAGGGCGAGGCGGAGCTGTGCCCACAGCAGCTGCAGCAGTTCTGCGCCGCATGCTTTCTGTCCCTGAGCATCAGTGTGTGTGCAGACGGGCAGCTGGACCGCCTCGCCCTGCGCATCCGGCCTGCCCGCCTCTACCTGGAGGACACCTTCATCTACTACATGAAGACGCTCTTCCACACCTACATTCCAGAGTGTGCAGCGGGGGGGCGGGGCTGTGTCAGGGGGGTGGAGTCAGCGGTGCCGCAGGAGGTGCTGGAGTCGATGCGTGCTCTGGTGTCGCCGCTGCGGCTGCAGAAACTGTCCATCGAGCCGCTGCAGCTGCTGGTCAGCATCCACGCCTCGCTGAAGCTCTACATCGCCTCTGACCACACGCCGCTGGCCTTCTCTGAGTTCGAGCGCGGGCCCGTCTGCACCACCGCACGACACCTGGTGCACACACTCGCCATGCACTACGCAGCAGGAGCACTCTTCAGGGCAGGCTGGGTTGTCGGCTCTCTGGAGATTCTGGGCAGTCCAGCCAGTCTGGTGCGTAGTATCGGGAACGGTGTGTCTGATTTCTTCCGTCTGCCGTATGAGGGTTTGACCCGCGGGCCAGGTGCGTTCATCAGCGGCGTCTCCAGAGGAACAAACTCCTTCATCAAACACATCTCTAAAGGAACGCTGACGTCCATCACGAATCTGGCCACGAGTCTGGCGCGTAACATGGACCGGCTGTCGCTGGACGATGAGCACTACACGCGGCAGGAGGAGTGGCGGCGGCAGCTGCCCGAAACACTGGGAGACGGACTGAGACAGGGGCTGTCCAGACTGGGCATCAGCCTGCTGGGAGCGGTCGCAGGAATCGTGGATCAGCCCATGCAGACGTTCACTCGTACGCTGGAGCTGCCAAACTCTGCGAGCAGCGCGGCCCGCGGCGTCATCTCCGGCGTGGGGAAAGGCATTGTGGGAGTTTTTACTAAACCAATCGGAGGAGCCGCTGAGCTGGTGTCACAGACGGGCTACGGCCTCCTCCATGGCGCAGGACTCTGGCAGCTCCCTAAACAGCTCCACCAGCCGACTGACAACCGATCCGCCGACGCCGCTAACAGCAGCGCCAAATACATCTGGAAGATGCTGCAGTCTCTGGGCCGTGCGGAGCTCCACATGGCGCTGGATGTGTGTATGGTGAGCGGCTCGGGGCAGGAACGCGCCGGCTGTTTGCTGCTGAGCGCTGAGGTGCTGTTTGTGGTCAGTGTGTGTGAGGACGCGCAGCAGCAGGCCTTCCCCATCACCGAGATCCAGTGTGAACACGACACACACACACCTGGACGCATCACACTCACACTGCAGCAGCAGCGCGTCAACACTGATGCTGAGGTCGAGGGCGTCCGCGAGCGTCTGTCGGAGCTGCAGTACACTCGGCTGCTGGATTTCGTGCGCGGTGCGTCTCCGTTTCTGCCCGCGGCGCTGCAGCCGAGCGTCAGTCCAGCAGAACCGCAGCGCAGCGTCTGCAGAACATACCAGTACCAGGCGGACGCCGCATTCGCACGCGTGTTCGTCTGCAGGTTCCAGATGGTGAAGAACAGAGCGCTGCGCATCGGCTTCCACTGACACACACACACACACACACACACACACACACACTCTGCTGCAGCACACATGGCCTTCTGTGTGTGTGTGTCTATAATGATGCCAAAAGAGACGATCCACTGTTTATATGTAATTTCCTGATTAAATTTGTGAGAAAT

